# Early metabolic reprogramming licenses *Streptococcus pneumoniae* for Influenza-driven superinfection

**DOI:** 10.64898/2026.01.10.698720

**Authors:** Marion Lagune, Niccolo Bianchi, Margaux Herren, Julien Prados, Aurelie Caillon, Grishma Kulkarni, Filomena Silva, Olivier Von Rohr, Roberto Sierra, Matthieu Colpaert, Matthew S Gentry, Jan-Willem Veening, Joachim Kloen, Mirco Schmolke, Simone Becattini

**Affiliations:** Department of Pathology and Immunology, Faculty of Medicine, University of Geneva, Geneva, Switzerland; Geneva Centre for Inflammation Research, University of Geneva, Geneva, Switzerland; Bioinformatics Support Platform for data analysis, Faculty of medicine, University of Geneva, Geneva, Switzerland; Department of Microbiology and Molecular Medicine, Faculty of Medicine, University of Geneva, Geneva, Switzerland; Department of Biochemistry and Molecular Biology, College of Medicine, University of Florida, USA; Department of Fundamental Microbiology, Faculty of Biology and Medicine, University of Lausanne, Lausanne, Switzerland

## Abstract

Bacterial pneumonia remains a major cause of morbidity and mortality following influenza A virus (IAV) infection. However, the adaptive mechanisms that enable pathogen expansion in the post-viral lung remain poorly defined. Here, using a mouse model of IAV–*Streptococcus pneumoniae* superinfection, we characterize bacterial transcriptional reprogramming *in vivo*. We identify alcohol dehydrogenases (AdhA and AdhE), that support NAD⁺ regeneration during mixed-acid fermentation, as key determinants of bacterial fitness specifically in the IAV-primed lung. Genetic deletion of these results in a pronounced fitness defect during superinfection but not in primary bacterial pneumonia. Consistent with this requirement, pharmacological inhibition of alcohol dehydrogenases limits bacterial expansion and dissemination following IAV infection. Mechanistically, we show that IAV infection profoundly remodels the lung environment, inducing hypoxia and increasing the availability of alternative carbon sources, which together impose a metabolic dependency on Adh for bacterial expansion. Our findings place metabolic adaptation as a central driver of pneumococcal outgrowth following viral infection and reveal exploitable vulnerabilities for therapeutic intervention.

## Introduction

Respiratory viral infections, including those caused by influenza A virus (IAV), are frequently complicated by subsequent bacterial infections, a phenomenon commonly referred to as superinfection, that can progress to severe pneumonia. Bacterial superinfections were a major cause of mortality during historical influenza pandemics^1–3^ and remain a critical clinical complication during seasonal and pandemic IAV outbreaks^4,5^. Although diverse bacterial species can cause superinfection, *Streptococcus pneumoniae* remains one of the leading etiological agents of severe post-influenza pneumonia ^4–6^.

The pathogenesis of viral–bacterial superinfection reflects a complex interplay between host immune responses^7^, viral factors^8–10^, bacterial virulence determinants^11,12^ and the resident respiratory microbiota^13^. During IAV infection, epithelial damage and disruption of lung homeostasis facilitate bacterial adhesion and invasion^14^, while virus-induced type I interferon responses impair antibacterial immunity^15^. In parallel, viral infection profoundly remodels the lung environment, creating conditions that influence bacterial physiology and growth. While host- and virus-driven mechanisms that promote superinfection have been extensively studied^7,16^, comparatively little is known about how bacterial pathogens adapt to the altered lung environment following viral infection. This gap is particularly striking given that, after IAV infection, *S. pneumoniae* can establish productive lung colonization from markedly reduced infectious doses^17,18^, suggesting that the bacterium engages distinct strategies to sense and exploit the virus-modified pulmonary niche. Defining these superinfection-specific bacterial programs is therefore essential to understand the mechanisms that support pneumococcal expansion in the post-viral lung and may reveal bacterial processes that can be therapeutically targeted to limit bacterial outgrowth. *In vivo* genetic screens performed in the context of superinfection have identified several bacterial determinants required for pneumonia^19–21^ ; however, these genes were also essential in primary pneumococcal pneumonia, suggesting that they represent core requirements for pneumococcal pathogenicity *in vivo* rather than adaptations specific to the lung environment shaped by prior viral infection. Bacterial responses that enable selective expansion during superinfection remain poorly defined. Here, we investigate bacterial adaptive responses to the virus-modified lung environment with dual-RNA sequencing in a mouse model of influenza A virus–*S. pneumoniae* superinfection. By adopting a bacterial-centric approach, we uncover metabolic strategies that specifically support pneumococcal expansion during superinfection. Our data provide mechanistic insight into post-viral pneumonia and highlight bacterial adaptive metabolic pathways as a potential druggable target to limit bacterial expansion following viral infection.

## Results

### Influenza-modified lung environment drives transcriptomic reprogramming in *S. pneumoniae*

Heightened susceptibility to secondary bacterial infection persists for weeks after IAV infection, despite complete viral clearance^22^. These findings suggest that *S. pneumoniae* detects and exploits persistent IAV-induced alterations in the lung microenvironment via a dedicated adaptation program. To identify bacterial genes required for *S. pneumoniae* adaptation to the IAV infection- shaped lung environment, we first established a temporal model of post-influenza susceptibility to pneumococcal superinfection (Fig. 1a). Mice were primed intranasally with PBS or a sublethal dose of IAV (Influenza A/Netherlands/602/2009)^23^. IAV-infected mice displayed transient weight loss, starting around day 7 post-infection, peaking at days 9–10, and recovering to baseline by days 13–14 (Extended Data Fig. 1a-c). We subsequently challenged both groups with *S. pneumoniae* (strain D39L derived from D39 background, serotype 2)^24^ 7, 14 or 28 days post priming, and sacrificed 24h later. Consistent with previous reports^17,18^, low dose bacterial challenge (∼10^4^ CFU) only established lung colonization after prior influenza infection (Fig. 1b). To account for the low colonization efficacy of *S. pneumoniae* without previous viral challenge and yield sufficient bacterial titers for transcriptomic analysis, we adjusted the bacterial inoculum in PBS-primed mice (∼10^8^ CFU). Heightened susceptibility to pneumococcal superinfection permitted substantial pneumococcal expansion in the lung at week 1 post viral challenge, which persisted up to week 2 (Fig. 1c). By week 4, PBS and IAV primed mice displayed comparable susceptibility to *S. pneumoniae* infections (Fig. 1d). Next, we performed transcriptomic analyses of *S. pneumoniae* at time points associated with superinfection susceptibility. Mice were primed with PBS or IAV and infected with *S. pneumoniae* one or two weeks later. Total RNA was extracted from the lungs 24 h after bacterial challenge. Quantitative PCR confirmed pneumococcal abundance across experimental groups (Extended Data Fig. 1d-e). High-depth sequencing (100–300 million reads per sample) enabled recovery of a substantial fraction of pneumococcal transcripts (mean = 469,510 reads, 0.37 % of total reads; mean coverage = 23 X; see details in Supplementary Table 1). Viral transcripts, which are known to persist long after clearance of infectious particles^25^ were detectable at week1 (0.08-0.42 % of total reads), but had virtually disappeared by week 2 (< 0.001 % of total reads, see details in Supplementary Table 1). Principal component analysis (PCA) revealed that *S. pneumoniae* adopts in the IAV-primed lungs a distinct transcriptional state from PBS-primed controls (Fig. 1e,f). We identified a core transcriptional response comprising 17 upregulated and 21 downregulated genes shared between weeks 1 and 2 (Fig. 1g and Extended Data Fig. 1f). In addition, a larger set of differentially expressed genes was uniquely detected at week 1, with fewer time point–specific changes at week 2. The expanded transcriptional response at week 1 may reflect the higher pneumococcal burdens observed at this time point and possibly underlies additional adaptation mechanisms operating early on after IAV infection.

**Figure 1.**
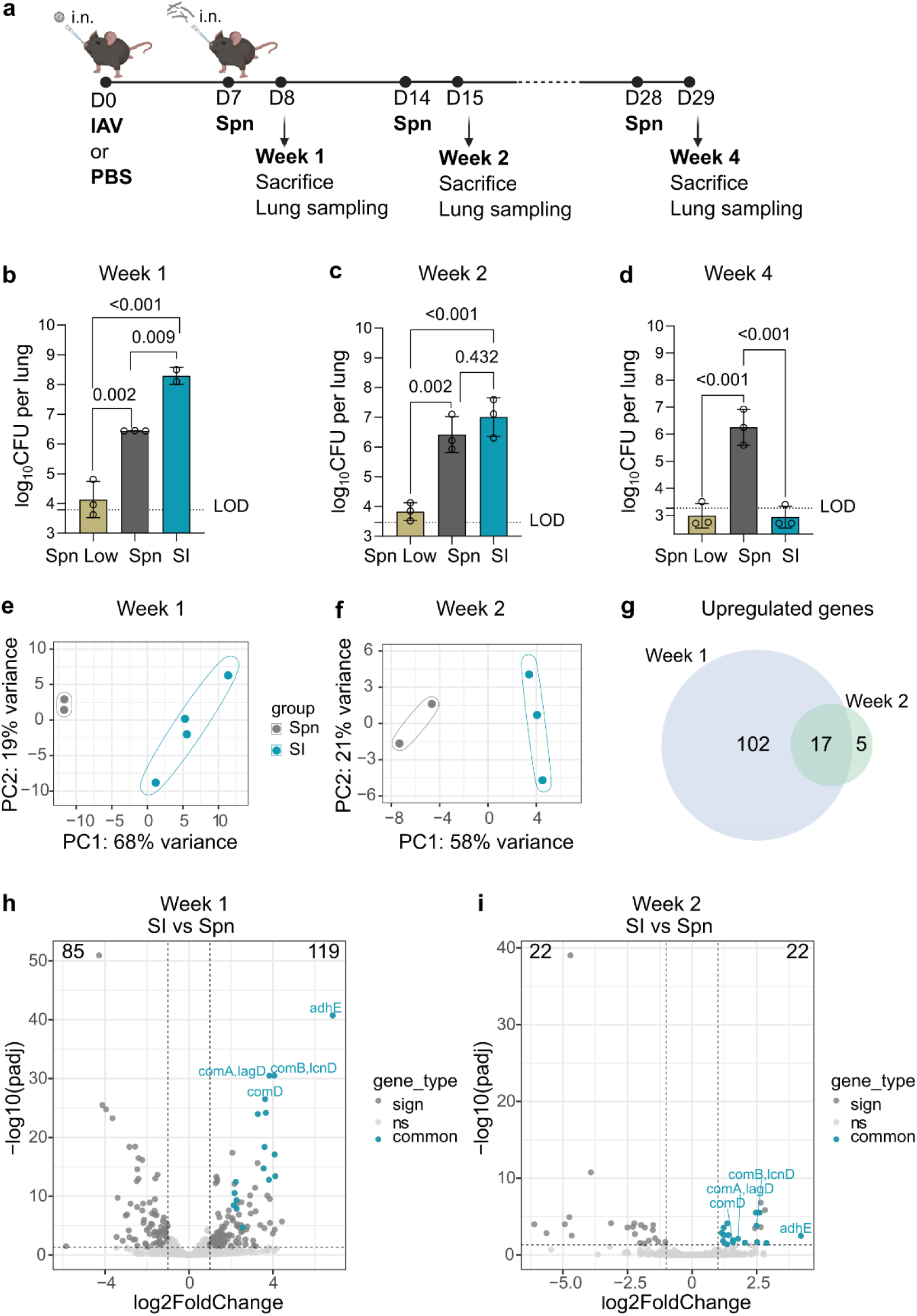
: Transcriptional adaptation of *S. pneumoniae* to the IAV-conditioned lung environment. **a**, Schematic of the mouse infection model. Mice were intranasally (i.n.) primed with PBS or influenza A virus (IAV; 10 PFU) and subsequently infected with *S. pneumoniae* D39L at 7, 14 or 28 d post IAV infection. Bacterial inocula were ∼1 × 10⁸ CFU for pneumococcal infection alone (Spn) and ∼1 × 10⁴ CFU for superinfected (SI) mice. Mice were euthanized 24 h after bacterial challenge. **b–d,** Lung bacterial burdens in mice infected with *S. pneumoniae* (**b**) 1 week, (**c**) 2 weeks or (**d**) 4 weeks following IAV infection. Dots indicate individual mice, bars depict geometric means ± geometric standard deviation (s.d.) (n=2-3, one representative of at least two experiments shown). Statistical significance was determined by ordinary one-way ANOVA with Tukey’s multiple-comparisons test. **e,f,** Principal component analysis (PCA) of *S. pneumoniae* transcriptomes recovered from Spn and SI mice 24h post inoculation, **(e)** 1 week and **(f)** 2 weeks post IAV infection. **g,** Venn diagram showing *S. pneumoniae* genes upregulated during SI at week 1 and week 2. **h,i**, Volcano plots showing differentially expressed *S. pneumoniae* genes during SI compared with Spn infection 24h post inoculation, **(h)** 1 week and **(i)** and 2 weeks post IAV infection. Genes commonly upregulated at both time points are highlighted in blue.

Gene set enrichment analysis revealed extensive transcriptional remodeling in pneumococcus, particularly at week 1 post-IAV infection, with quorum sensing representing the only pathway commonly modulated across both time points (Extended Data Fig. 1g,h). Among the downregulated gene sets we found fatty acid metabolism and biosynthetic pathways at week 1 post-IAV infection. We focused subsequent analyses on commonly upregulated genes, assuming that increased expression is more likely to reflect active bacterial adaptation to environmental cues. At both time points, we observed consistent induction of competence-associated genes and alcohol dehydrogenases, suggesting that these functions may contribute to *S. pneumoniae* adaptation to the IAV-altered lung environment (Fig. 1h,i). Collectively, these findings indicate that *S. pneumoniae* engages a dynamic and context-dependent transcriptional program in response to lung remodeling induced by prior viral infection. These programs potentially sustain bacterial fitness over the extended window of susceptibility to superinfection.

### Alcohol dehydrogenases are key determinants of *S. pneumoniae* adaptation and fitness during superinfection

To causally link the RNA-seq–identified genes to pneumococcal fitness during superinfection, we generated isogenic deletion mutants targeting genes of interest and evaluated their fitness *in vivo*. We first examined components of the competence system, including the *comAB* competence peptide transporter and the *comD* competence receptor. In competitive infection assays, deletion of *comD* did not produce a detectable fitness defect, whereas the *comAB* mutant exhibited a modest growth disadvantage during superinfection (Extended Data Fig. 2a,b). However, when mice were infected with either the wild-type or *comAB* mutant strains alone, in the presence or absence of prior IAV infection, no significant differences in lung bacterial burden were observed (Extended Data Fig. 2c). Thus, our data indicate that the competence system does not substantially contribute to pneumococcal superinfection.

We next focused on two genes encoding alcohol dehydrogenases, *adhE* and *adhA*. Both enzymes contribute to regeneration of the NAD⁺ pool during fermentative metabolism ^26^ (Extended Data Fig. 3a). *AdhE* is a bifunctional enzyme with both alcohol and acetaldehyde dehydrogenase activities ^27^ and was among the most highly upregulated genes at both week 1 and week 2 post-IAV infection, whereas *adhA* induction was restricted at week 1. Fitness assays of the corresponding single deletion mutants revealed no detectable defect for either strain *in vivo* (Fig. 2a,b). Since these enzymes operate within the same metabolic pathway and may serve overlapping functions, we assessed compensatory expression by qPCR. While *adhA* expression was unchanged in the *adhE* deficient strain, *adhE* expression was strongly upregulated in the *adhA* deficient strain (Fig. 2c,d), indicating possible functional redundancy.

**Figure 2:**
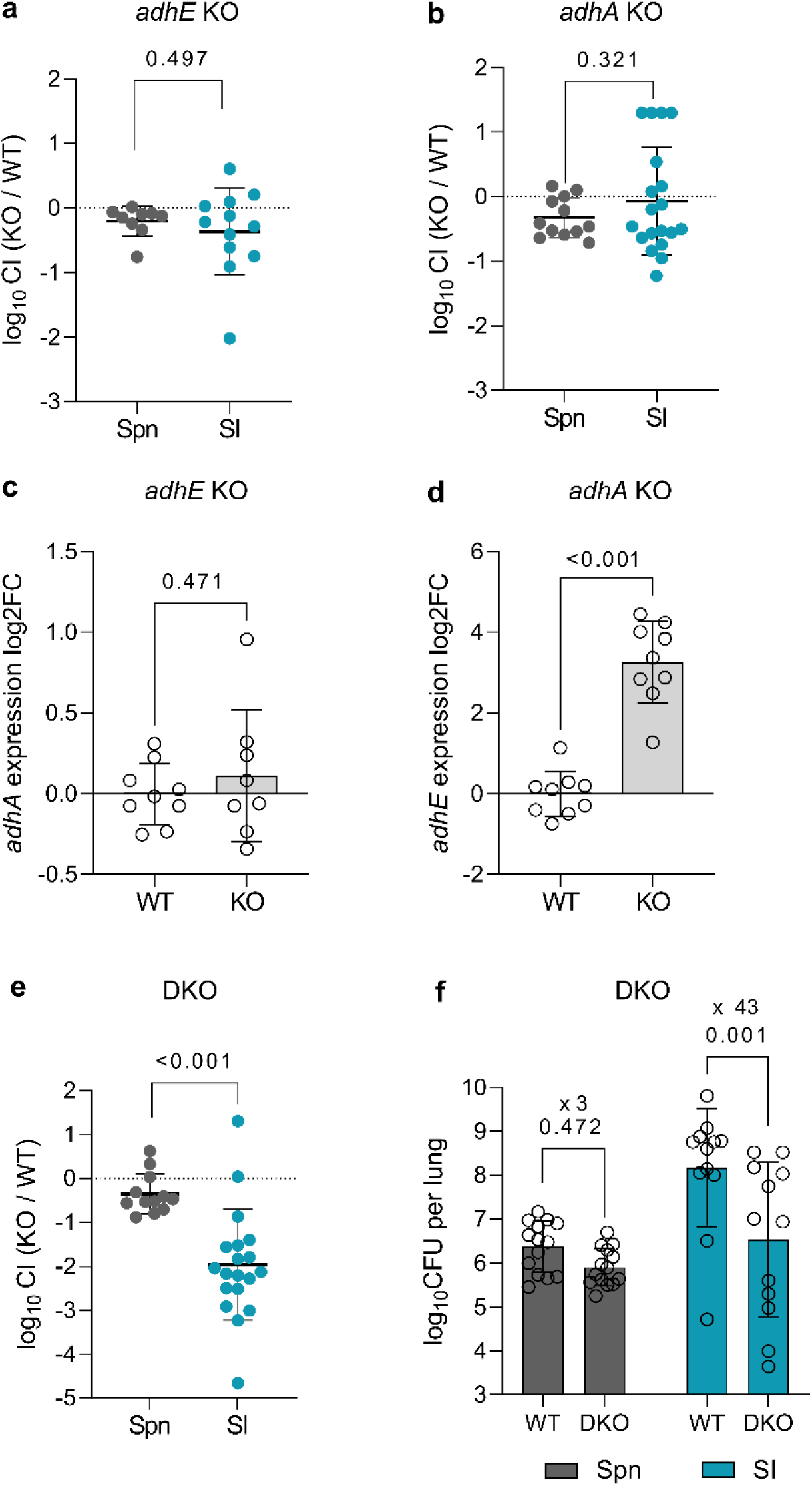
Alcohol dehydrogenases are required for *S. pneumoniae* adaptation during superinfection. **a**,**b**, Competitive infection with WT and isogenic deletion mutant **(a)** *adhE* KO and **(b)** *adhA* KO. Mice primed with PBS or IAV one week prior were infected with a 1:1 ratio of KO to WT bacteria, and sacrificed 24h later to determine relative burden. Dots represent individual mice (**a**, n= 9-12 per group; **b,** n= 12-19 per group; data pooled from 2 independent experiments; geometric means ± geometric s.d. are shown). **c,d**, Expression of **(c)** *adhA* and **(d)** *adhE* in the *adhE* KO and *adhA* KO strains, respectively, as determined by RT-qPCR, normalized to housekeeping gene in WT. Dots represent individual bacterial cultures (n = 8-9, data pooled from 3 independent experiments; means ± s.d. are shown). **e**, Competitive infection with WT and *adhE/adhA* DKO strain, performed as in (**a**,**b**). Dots represent individual mice (n = 12-19 per group, data pooled from 2 independent experiments; geometric means ± geometric s.d. are shown). **f**, Bacterial burdens 24h post individual infection with WT or *adhE/adhA* DKO strains in mice primed with PBS (Spn) or IAV (SI) 7 days prior. Dots represent individual mice (n= 14-12 per group, data pooled from 2 independent experiments; geometric means ± geometric s.d. are shown). Statistical significance was determined by two-tailed unpaired *t*-tests**(a,b,c,d,e)** and ordinary two-way ANOVA with Šídák’s multiple-comparisons test **(f).**

To test this directly, we generated a double knock-out (DKO) strain lacking both *adhE* and *adhA* (Extended Data Fig. 3b-c). The DKO grew normally *in vitro* (Extended Data Fig. 3d), excluding any intrinsic fitness defect. Remarkably, in competitive infection assays *in vivo*, the DKO was specifically outcompeted by the wild-type strain during superinfection, while displaying no fitness defect during primary pneumococcal pneumonia (Fig. 2e). We further performed single-strain infections to assess intrinsic fitness independently of competition with the wild-type strain. Although the DKO showed only a mild reduction in bacterial burden during primary infection, in superinfected mice it exhibited a ∼40-fold decrease in CFU relative to the wild-type strain (Fig. 2f). Together, these results demonstrate that AdhE and AdhA act redundantly and are jointly required as key metabolic determinants of bacterial fitness for optimal pneumococcal adaptation to the IAV-altered lung.

### IAV-induced lung remodeling triggers early *adhE* induction in *S. pneumoniae*

Genes encoding alcohol dehydrogenases were upregulated in the bacterial transcriptome at 24 h post-infection, a time point at which *S. pneumoniae* has already undergone substantial expansion in the IAV-primed lung. This transcriptional reprogramming could therefore reflect either an early adaptive response to the IAV-modified environment, or a secondary metabolic consequence of bacterial outgrowth at later stages of infection. To discriminate between these two possibilities, we performed an *in vivo* time-course experiment to monitor pneumococcal growth during the first six hours following bacterial challenge. *S. pneumoniae* expanded substantially within two hours in IAV-primed mice, whereas bacterial titers declined over the same period in PBS-primed mice infected with a comparable dose (Spn Low) (Extended Data Fig. 4a,b). These findings indicate that the IAV-primed lung environment promotes early pneumococcal adaptation, likely independent of bacteria-driven host remodeling. We next asked whether alcohol dehydrogenase up-regulation is part of such early adaptive response. To ensure sufficient bacterial biomass in the respiratory tract for reliable detection of pneumococcal transcripts by RNA-seq, IAV- or PBS-primed mice were infected with the higher inoculum previously used to induce pneumococcal pneumonia in PBS-treated animals, and sacrificed two hours later (Fig. 3a-b). PCA revealed that *S. pneumoniae* adopts a distinct transcriptional profile in the IAV-primed lung as early as 2 hours post-infection (Fig. 3c) although, as expected at this early time point, relatively few genes were differentially expressed (Fig. 3d). Notably, *adhE* emerged among the most strongly upregulated transcripts and was the only gene consistently induced both at this early stage and at 24 h following superinfection at weeks 1 and 2 post-IAV infection (Fig 3d and Extended Data Fig. 4c). Together, these results highlight not only a central role for AdhE in promoting pneumococcal fitness during superinfection, but also indicate that its induction is rapidly triggered by local factors encountered in the lung tissue as a consequence of IAV infection.

**Figure 3:**
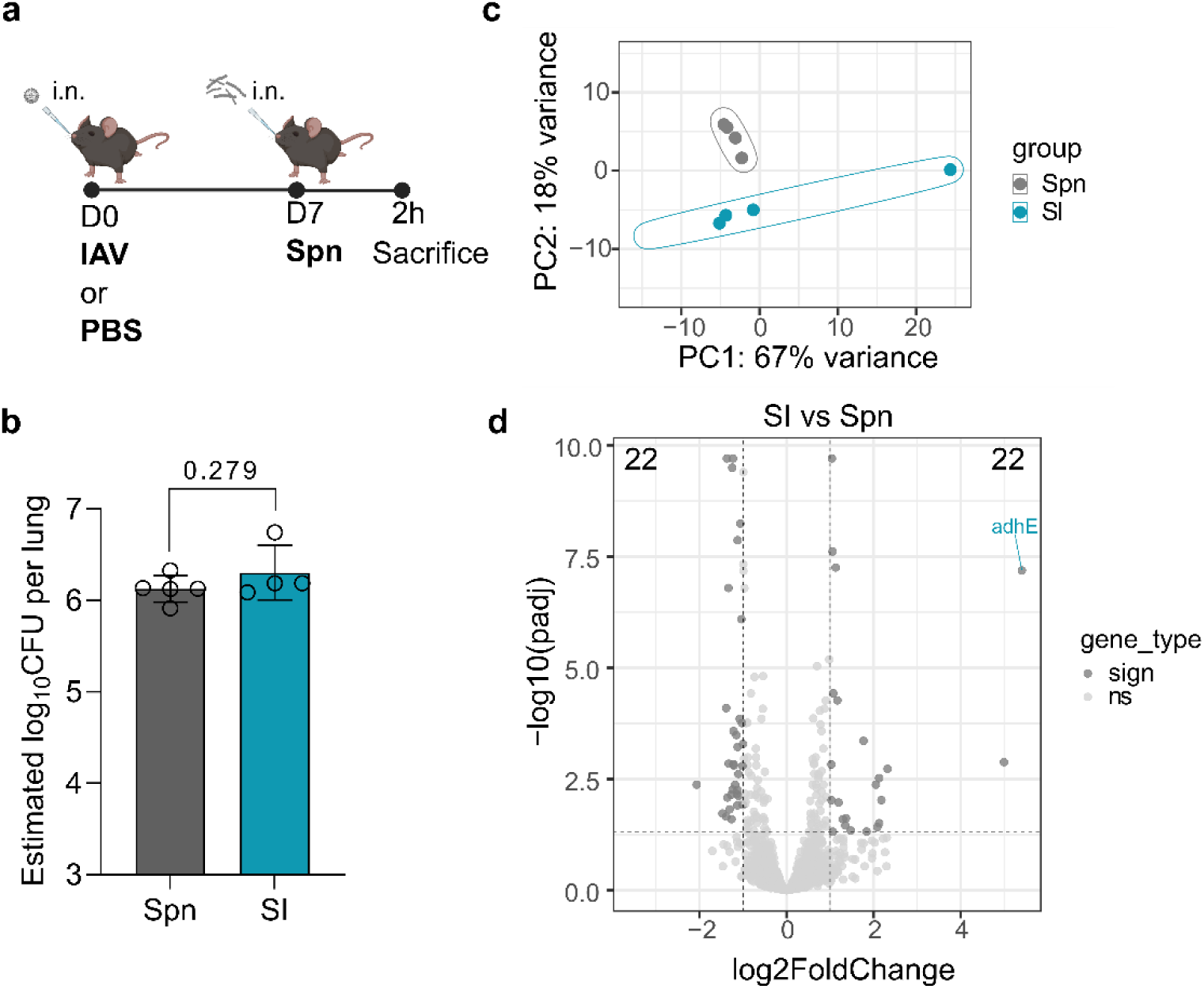
IAV-modified environment promotes early bacterial adaptation. **a**, Schematic of the mouse infection model. Mice were intranasally (i.n.) primed with PBS or influenza A virus (IAV; 10 PFU) and subsequently infected with *S. pneumoniae* D39L at 7d post IAV infection. Bacterial inoculum was ∼1 × 10⁸ CFU for both pneumococcal infection alone (Spn) and superinfected (SI) mice. Mice were euthanized 2 h after bacterial challenge. **b**, Estimation of bacterial titer based on expression of the housekeeping gene *gyrB,* as determined by RT-qPCR. Dots indicate individual mice (n = 4-5; geometric means ± geometric s.d. are shown). Statistical significance was determined by two-tailed unpaired *t*-tests. **c**, Principal component analysis (PCA) of *S. pneumoniae* transcriptomes recovered from Spn and SI mice, infected as explained in **(a)**, from lungs harvested 2 h post infection. **d**, Volcano plots depicting differentially expressed *S. pneumoniae* genes during SI compared with Spn infection at 2 h post infection. Non-significant genes are shown in light grey, and significant genes (|L2FC| > 1, padj < 0.05) are shown in dark grey. *adhE* is highlighted in blue.

### Immune-associated changes in the lung milieu during superinfection susceptibility

As *adhE* is crucial to establish superinfection, the above results prompted us to interrogate host-derived cues that may be responsible for its induction. IAV infection elicits a robust inflammatory response in the respiratory tract, characterized by the recruitment of innate and adaptive immune cells from the circulation^28^, the production of inflammatory mediators^29^ and the extensive remodeling of the pulmonary metabolite landscape^30^. While these responses are essential to viral clearance, they come at the cost of compromised tissue integrity and collateral lung damage^31^. In fact, in severe pneumonia the accumulation of immune cells and fluid within the airspaces hinders gas exchange and is a hallmark of poor clinical outcome^32^.

Flow cytometry analysis at week 1 post-IAV infection confirmed a marked remodeling of the pulmonary immune compartment, with a reduction in alveolar macrophages as described previously^33^ and a pronounced increase in multiple myeloid populations, including interstitial macrophages, monocytes, eosinophils, neutrophils, monocyte-derived and classical dendritic cells (Extended Data Fig. 5). Lymphoid populations including B cells, natural killer cells, CD4+ and CD8+ T cells, and γδ T cells also accumulated in the lung tissue (Extended Data Fig. 6).

We next analyzed host transcriptomic changes by comparing lung gene expression at weeks 1, 2, and 4 post-IAV infection, reasoning that local host factors promoting superinfection would remain altered at weeks 1 and 2 but return to baseline by week 4, when susceptibility is lost (Extended Data Fig. 7a). Principal component analysis indicated that transcriptomes from IAV-infected mice were clearly separated from mock-infected controls at week 1 along PC1, and that this difference progressively diminished, returning toward control levels by weeks 2 and 4 (Extended Data Fig. 7b), in agreement with published data^34^. Consistent with this observation, infected lungs exhibited the most extensive transcriptional remodeling and largest number of upregulated genes at week 1 (Extended Data Fig. 7c). A total of 394 host genes were upregulated at both weeks 1 and 2, when susceptibility to superinfection persists, but returned to baseline at week 4, fulfilling our criteria for local host factors potentially contributing to *S. pneumoniae* adaptation. Gene set enrichment analysis indicated robust upregulation of multiple immune-related pathways, in line with our flow cytometry analyses (Extended Data Fig. 7d). At the gene level, cytokines and cytokine receptors, including *Ifnγ*, *Il21* and *Il21r,* were strongly upregulated following IAV infection (Extended Data Fig. 7e). IFN-γ signaling has previously been shown to promote *S. pneumoniae* expansion following IAV infection, at least in part by impairing alveolar macrophage phagocytic capacity, but whether this impacts pneumococcal metabolism is not known^35^. Consistent with the above study, IFN-γ blockade reduced bacterial burden during superinfection, however it did not alter *adhE* expression in *S. pneumoniae* (Extended Data Fig. 7f-h). Thus, although IFN-γ promotes bacterial outgrowth, it does so independently of *adhE* regulation, indicating that restoration of NAD⁺ is not a contributing mechanism. We next applied a similar approach to investigate IL21 and IL-21R, whose role in superinfection has not been explored previously (Extended Data Fig. 7i). IL-21R blockade did not affect bacterial titers, nor did it modulate *adhE/A* expression, suggesting that IL-21 signaling does not contribute to bacterial expansion (Extended Data Fig. 7j,k). Together, these results demonstrate that IAV infection induces extensive and long-lasting remodeling of the host pulmonary environment, including a core set of upregulated host immune genes. While some of these factors directly promote bacterial outgrowth, others likely reflect collateral immune activation. Importantly, none of these immune mediators explained the *adhE* dependency observed during *S. pneumoniae* superinfection, pointing instead to alternative host-derived drivers of this phenotype.

### Hypoxia and a shift towards alternative carbon sources induce alcohol dehydrogenases in *S. pneumoniae*

During glycolysis, bacteria reduce NAD⁺ to NADH; to replenish the limited intracellular NAD⁺ pool and sustain glycolytic carbon flux, swift re-oxidation of NADH is essential^36^. Alcohol dehydrogenases function at a key branch point in pyruvate metabolism, influencing both carbon flux and cellular redox balance. Under standard laboratory conditions, i.e. growth in glucose-rich medium and in the presence of oxygen, *S. pneumoniae* preferentially relies on lactate dehydrogenase (Ldh) to convert pyruvate into lactate and regenerate NAD⁺ following glycolysis^37^ (Extended Data Fig 3a). However, several *in vitro* studies have shown that *S. pneumoniae* can shift toward mixed-acid fermentation producing formate, acetate, and ethanol, particularly when alternative sugars such as galactose are present^38^. We reasoned that immune cell infiltration and inflammation-driven tissue remodeling, including increased vascular leakage into the alveolar space^39^, may reshape the local carbon landscape, potentially impacting the mechanism of NAD⁺ regeneration.

Although global carbohydrate metabolism in *S. pneumoniae* was not significantly altered based on gene set enrichment analysis, we observed upregulation of multiple sugar metabolism–related genes at week 1 post-IAV infection (Extended Data Fig. 8a). These included several genes encoding carbohydrate transporters with both narrow and broad substrate specificity. Notably, genes involved in the uptake and utilization of maltodextrin, a mixture of glucose oligomers (3-20 units linked by α(1→4) glycosidic bonds), including those belonging to the *malXCD* and *malMP* operons, were among the most strongly induced (Extended Data Fig. 8a). These observations are consistent with IAV infection altering carbohydrate availability in the lung environment. To test this, we performed untargeted GC-MS-based metabolomic profiling on cell pellets from BALF of mock-treated and IAV-infected mice. This analysis revealed a significant increase in maltose and mannose in samples from IAV-infected mice, whereas other measured carbohydrates showed no major difference between conditions (Fig. 4a, b and Extended Data Fig. 8b-e). We next asked whether these sugars affect expression of *adhE. S. pneumoniae* was briefly cultured in chemically defined medium supplemented with glucose, maltose, or mannose, followed by targeted qRT-PCR analyses. Both maltose and mannose significantly induced *adhE* expression compared with glucose (Fig. 4c). We further tested the effect of maltotriose and glycogen, a glucose polymer that *S. pneumoniae* can hydrolyze into maltodextrin^40^. Strikingly, both substrates induced *adhE* expression (Extended Data Fig. 8f). This phenotype extended to other sugars, including galactose as well as the disaccharides sucrose, trehalose, and isomaltose (Extended Data Fig. 8g). Similar, but generally more modest patterns of upregulation were observed for *adhA*, although this was not induced by mannose or isomaltose, indicating partial overlap in the carbon sources that can promote expression of distinct alcohol dehydrogenases (Extended Data Fig. 8h, i). Together, these results indicate that exposure to multiple non-glucose carbon sources present in the IAV-infected lung, including maltose/maltodextrin, is sufficient to drive *adhE* upregulation in *S. pneumoniae*.

**Figure 4:**
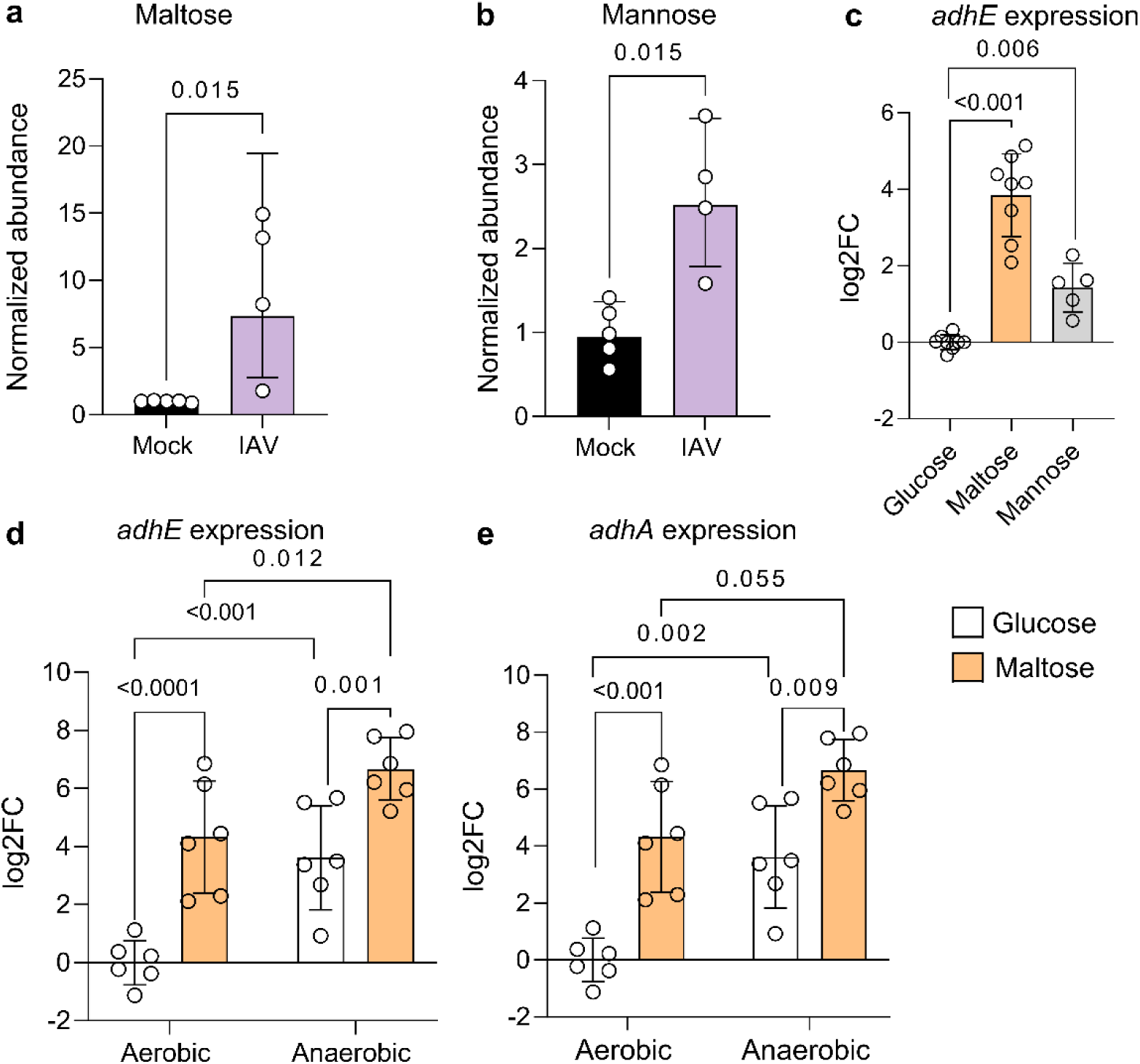
IAV infection reshapes lung oxygen and carbon availability to regulate pneumococcal *adh* expression. **a,b** Maltose **(a)** and mannose **(b)** levels in cells from bronchoalveolar lavage fluid from mock- or IAV-infected mice, measured by targeted metabolomic analysis using GC–MS. Metabolite levels are shown as relative abundance normalized to mock-infected samples and to total sample signal. Dots indicate individual mice, bars depict geometric means ± geometric s.d. (n = 4-5; one representative of at least two experiments shown). **c**, *AdhE* expression in *S. pneumoniae* cultured in chemically defined medium supplemented with glucose, maltose or mannose for 30 min. Dots represent individual bacterial cultures (n = 5-8, data pooled from at least two independent experiments; means ± s.d. are shown). **d,e**, *adhE* **(d)** and *adhA* **(e)** expression in *S. pneumoniae* cultured in chemically defined medium supplemented with glucose or maltose under aerobic and anaerobic conditions for 30 min. Dots represent individual bacterial cultures (n = 6, data pooled from at least two independent experiments; means ± s.d. are shown). Statistical significance was determined by unpaired two-tailed Mann–Whitney tests **(a,b)**, two-tailed unpaired *t*-tests **(c)** and 2-way ANOVA **(d,e).**

Under aerobic conditions, NAD⁺ regeneration in pneumococcus relies primarily on the oxygen-dependent NADH oxidase Nox and the pyruvate oxidase SpxB, the latter limiting lactate production^41,42^. In our samples, *spxB* was downregulated in the pneumococcal transcriptome at weeks 1 and 2 post–IAV infection (Extended Data Fig. 3a; Extended Data Fig. 9a-b), suggesting reduced oxygen availability in the tissue. This interpretation aligns with prior reports of IAV-induced hypoxia in the lung parenchyma, as well as with the upregulation of hypoxia-responsive host genes, including *Hif1a*, one week post-IAV infection (Extended Data Fig. 9c)^43^. Further corroborating this possibility, other central hypoxia-responsive genes were upregulated in this dataset (Extended Data Fig. 9c). These findings suggest that oxygen-dependent NAD⁺ regeneration may be compromised *in vivo* during IAV infection, possibly prompting reliance on alternative pathways. To directly assess the impact of oxygen availability on *adh* genes expression, we cultured *S. pneumoniae* under aerobic and anaerobic conditions in the presence of either glucose or maltose. Both *adhE* and *adhA* were potently induced under anaerobic conditions, with maltose consistently eliciting a stronger response than glucose (Fig. 4d,e). Taken together, these data suggest that IAV infection reshapes the lung microenvironment by simultaneously altering carbon source availability and reducing oxygen tension, with these cues acting synergistically to induce *adh* expression in *S. pneumoniae*.

### Alcohol dehydrogenases are upregulated across serotypes in superinfection

*S. pneumoniae* displays extensive genetic and phenotypic diversity, with serotype- and strain-specific differences influencing host colonization, virulence, and environmental adaptation^44,45^. To assess whether our findings on the adaptation of pneumococcus to the IAV-primed lung extend beyond the D39L strain (serotype 2), we performed parallel experiments using the strain ATCC-6303, a serotype 3 isolate. Owing to its increased virulence, PBS-primed mice were infected at day 7 post-priming with 10^6^ and 10^3^ CFU, and IAV-primed mice were infected at day 7 post-priming with 10^3^ CFU. Similar to D39L, ATCC-6303 could exploit a prolonged window of susceptibility in the IAV-modified lung environment (Fig. 5a–c). Having confirmed comparable pneumococcal burdens across conditions by qPCR (Extended Data Fig. 10a), we performed transcriptomic analyses of whole-lung RNA. Principal component analysis revealed that ATCC-6303, like D39L, adopts a transcriptional state in the IAV-primed lung that is distinct from PBS-primed controls at both week 1 and week 2 post-infection (Fig. 5d).

**Figure 5:**
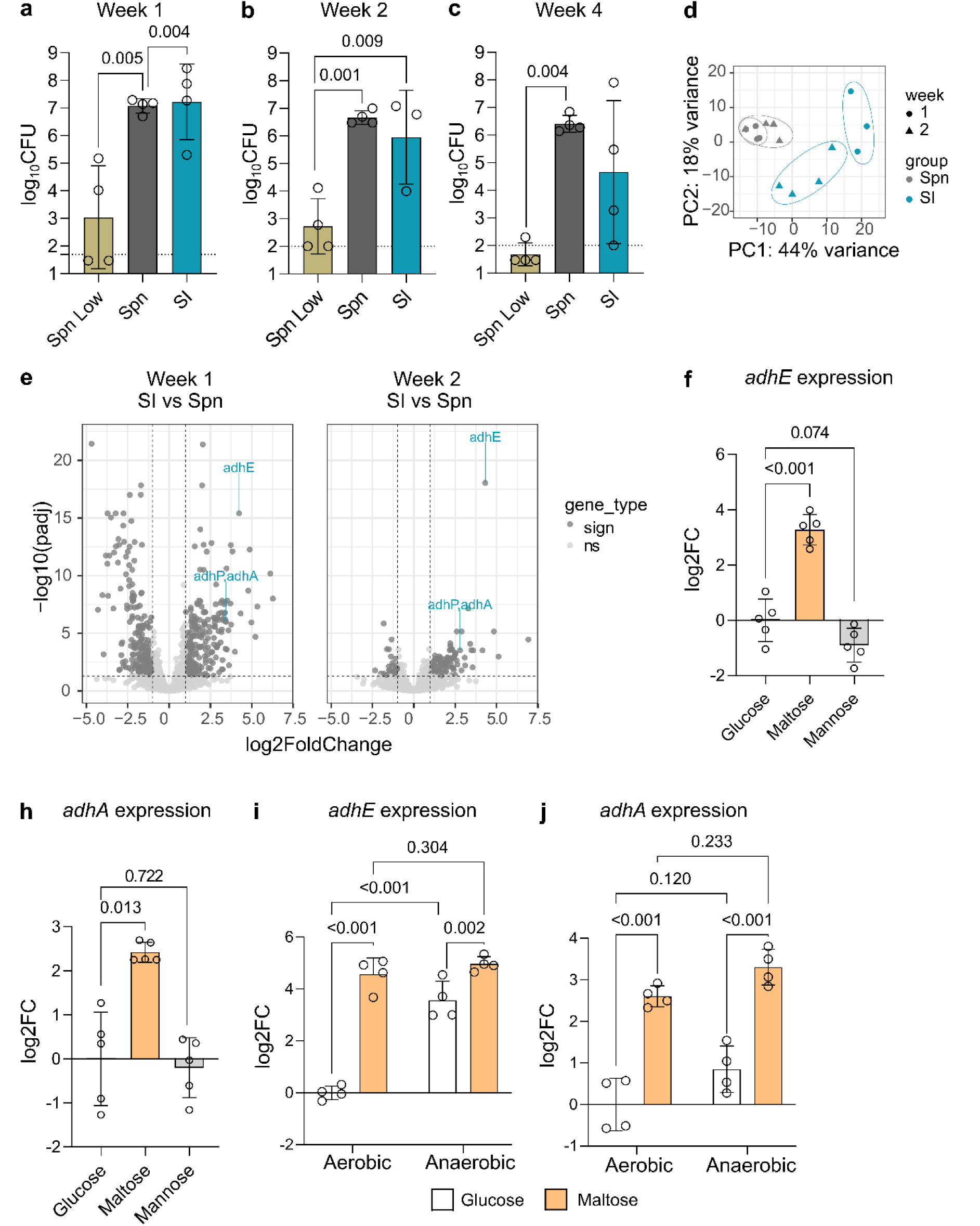
Regulation of *adhA* and *adhE* expression upon superinfection is conserved in a serotype 3 strain ATCC-6303. **a–c**, Lung bacterial burdens in mice infected for 24h with *S. pneumoniae* at **(a)** 1 week, **(b)** 2 weeks or **(c)** 4 weeks following IAV priming. Dots indicate individual mice, bars depict geometric means ± geometric standard deviation (s.d.) (n=3-4, one representative of at least two experiments shown). **d,** Principal component analysis (PCA) of *S. pneumoniae* transcriptomes recovered from Spn and SI mice 24h post infection, 1 week (dots) and 2 weeks (triangles) post IAV priming. **e**, Volcano plots showing differentially expressed *S. pneumoniae* genes during SI compared with Spn infection at week 1 (left) and week 2 (right). Non-significant genes are shown in light grey, and significant genes (|L2FC| > 1, padj < 0.05) are shown in dark grey. *adhE* and *adhA* are highlighted in blue. **f,h**, *adhE* **(f)** and *adhA* **(h)** expression in *S. pneumoniae* cultured in chemically defined medium supplemented with glucose, maltose or mannose. Dots represent individual bacterial cultures (n = 5, data pooled from at least two independent experiments; means ± s.d. are shown). **i,j**, *adhE* **(i)** and *adhA* **(j)** expression in *S. pneumoniae* cultured in chemically defined medium supplemented with glucose or maltose under aerobic or anaerobic conditions. Dots represent individual bacterial cultures (n = 4, data pooled from at least two independent experiments; means ± s.d. are shown). Statistical significance was determined by ordinary one-way ANOVA with Tukey’s multiple-comparisons test **(a-c)**, two-tailed unpaired *t*-tests **(f,h)** and two-way ANOVA with Fisher’s LSD test **(i,j)**.

Comparative analysis revealed a larger number of differentially expressed genes in ATCC-6303 than in D39L (Extended Data Fig. 10 b, c). A conserved core of 60 upregulated genes was shared between strains and included among the top hits *adhE* and *adhA*, competence-associated genes, and iron uptake systems (Fig. 5e and Extended Data Fig. 10d, e). Furthermore, genes involved in carbohydrate uptake and metabolism were also induced in ATCC-6303, including components of the maltodextrin utilization e.g. *malP*, *malG* and *malX* (Extended Data Fig. 10f). Strain-specific transcriptional responses were also observed; for example, genes involved in arginine biosynthesis (*argG* and *argH*) were selectively upregulated in ATCC-6303 (Extended Data Fig. 10 d, e ).

Based on the conservation of *adhE/A* and *malP/G/X* upregulation in the ATCC-6303 strain during superinfection, we next assessed the importance of alternative carbon sources in transcriptional reprogramming, and cultured the bacterium in the presence of various mono-, di-, tri- and polysaccharides. Similar to what observed for D39L, maltose and polysaccharide sources of maltose robustly induced both *adhE* and *adhA* (Fig. 5f,h and Extended Data Fig. 11a-d), whereas galactose and sucrose elicited weaker induction (Extended Data Fig. 10a, b). In contrast to D39L, mannose and arabinose had no effect (Fig. 5f,h and Extended Data Fig. 11a, b). We further examined the contribution of oxygen availability to *adh* regulation. Anaerobic growth led to increased *adhE* expression in presence of glucose compared with aerobic conditions, whereas growth in maltose resulted in similar *adhE* upregulation irrespective of oxygen availability (Fig. 5i,j). In contrast, *adhA* expression was largely insensitive to oxygen tension under both carbon sources. Importantly, ATCC-6303 retained the ability to sense oxygen availability, as indicated by downregulation of *spxB* under anaerobic conditions (Extended Data Fig. 11e). Together, these data suggest that in the ATCC-6303 strain, carbon source availability exerts a dominant influence over *adh* induction, with hypoxia playing a more limited modulatory role compared to observed in strain D39L. We conclude that induction of alcohol dehydrogenases represents a conserved adaptive response of *S. pneumoniae* to the IAV-modified lung environment across serotypes. At the same time, the environmental cues driving this response, particularly carbon source utilization and sensitivity to hypoxia, are differentially integrated depending on strain background.

### Pharmacological targeting of ADH restrains superinfection

The strong dependence of pneumococcal superinfection on alcohol dehydrogenase suggested an attractive therapeutic opportunity. To explore whether this metabolic vulnerability could be pharmacologically exploited, we used fomepizole, an FDA-approved broad-spectrum inhibitor of alcohol dehydrogenases used to treat methanol poisoning and previously applied in models of pneumococcal lung infection to modulate antibiotic sensitivity^26^ (Fig. 6a). Strikingly, in the context of superinfection, mice treated with fomepizole displayed significantly lower bacterial loads in the lung compared with untreated superinfected controls (Fig. 6b). By contrast, during primary pneumococcal infection, fomepizole provided no benefit and instead led to a modest increase in pulmonary bacterial titers. These results indicate that the therapeutic effect of alcohol dehydrogenase inhibition is specific to the superinfection setting. Superinfected mice treated with fomepizole also displayed reduced bacterial dissemination, with fewer bacteria detected in the spleen, likely reflecting the diminished bacterial burden in the lungs (Fig. 6c). Treatment of ATCC-6303 superinfected mice with fomepizole significantly reduced bacterial burden in the lung, although to a lower extent than what observed for D39L, without impacting systemic spread (Fig. 6d,e). These findings indicate that alcohol dehydrogenase activity contributes to ATCC-6303 fitness during superinfection, albeit to a lesser extent than in the D39 background.

**Figure 6:**
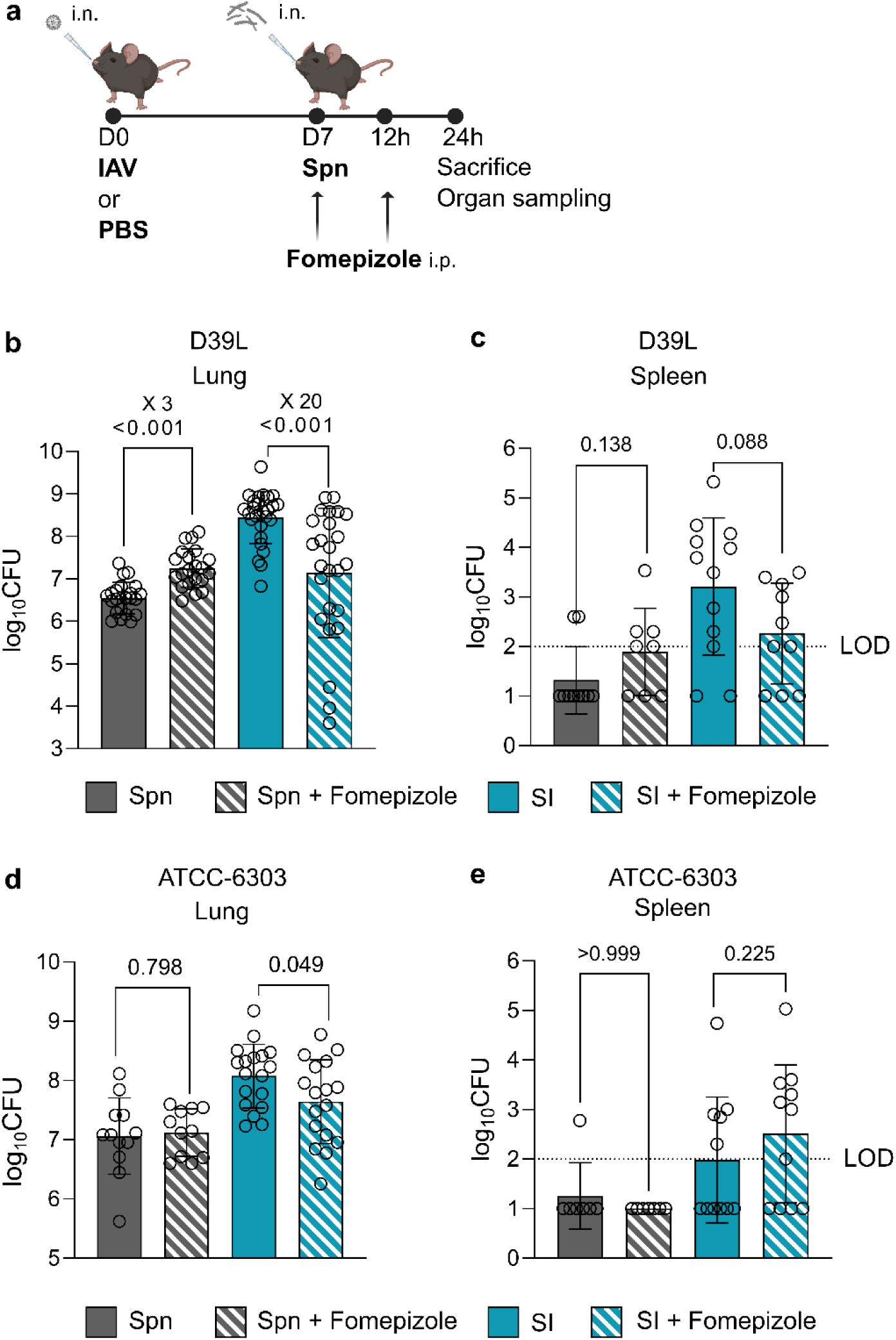
Pharmacological Inhibition of *S. pneumoniae* alcohol dehydrogenase limits bacterial expansion during superinfection. **a**, Schematic of the mouse infection model. Mice were intranasally (i.n.) primed with PBS or influenza A virus (IAV; 10 PFU) and one week later infected with *S. pneumoniae* ATCC-6303. Bacterial inocula were ∼1 × 10⁸ CFU (D39L) or ∼1 × 10⁶ CFU (ATCC-6303) for pneumococcal infection alone (Spn) and ∼1 × 10⁴ CFU (D39L) or ∼1 × 10³ CFU (ATCC-6303) for superinfected (SI) mice. Fomepizole (10 mg kg⁻¹ in 200 ul of PBS) or PBS were administered intraperitoneally at the time of bacterial infection and again 12 h post infection. Mice were euthanized 24 h after bacterial challenge. **b,c**, Lung **(b)** and spleen **(c)** bacterial burdens in mice infected with *S. pneumoniae* D39L. **d,e**, Lung **(d)** and spleen **(e)** bacterial burdens in mice infected with *S. pneumoniae* ATCC-6303. Data are pooled from at least two independent experiments. Dots indicate individual mice (lungs n = 12-26, spleens n = 6-12), bars represent geometric means with geometric s.d. **(b-e)**. Statistical significance was determined by two-tailed unpaired *t*-tests **(b-e)**.

Overall, pharmacological inhibition of alcohol dehydrogenase limits pneumococcal expansion specifically in the context of IAV superinfection, mirroring the context-dependent phenotype observed in the DKO strain.

## Discussion

Influenza A virus infection induces extensive^31^ and long-lasting remodeling of the lung environment^22^, generating a permissive niche for pneumococcal expansion that persists beyond viral clearance. While host immune dysfunction, epithelial damage, and viral virulence factors are well-established drivers of superinfection^46^, how bacterial pathogens actively sense and exploit the post-viral lung environment has remained comparatively underexplored. Here, by applying a bacterial-centric *in vivo* transcriptomic approach across multiple time points and pneumococcal strain backgrounds, we identify a dedicated metabolic adaptation program that enables *S. pneumoniae* to expand specifically in the influenza-altered lung. Our data place bacterial redox metabolism, and in particular alcohol dehydrogenase–mediated NAD⁺ regeneration, at the center of pneumococcal fitness during superinfection.

A key conceptual advance of this study is the demonstration that pneumococcal adaptation to the IAV-primed lung is both rapid and sustained. Transcriptional reprogramming was detectable as early as two hours after bacterial challenge, preceding substantial bacterial expansion, and was inducible for up to two weeks following viral infection. This timing argues strongly against a model in which bacterial gene expression changes are merely a secondary consequence of inflammation driven by bacterial overgrowth. Instead, pneumococcus appears to directly sense virus-induced environmental cues and engage an anticipatory metabolic program that licenses expansion in an otherwise hostile niche. This finding complements prior work describing host-driven permissiveness during superinfection^47^ and adds an active bacterial dimension to the pathogenesis of post-influenza pneumonia.

The superinfection-associated transcriptional program comprises multiple functional modules. Notably, genes related to competence and quorum sensing, iron acquisition, and carbohydrate transport were upregulated during superinfection, although the relative contribution of each axis to superinfection biology remains to be determined. While we show that competence genes are dispensable for *S. pneumoniae* expansion and superinfection *per se*, their induction may nonetheless facilitate horizontal DNA acquisition, with potential consequences for the emergence of antibiotic resistance. In addition, the downregulation of fatty acid biosynthesis observed in this system mirrors immune-driven adaptations reported for other Gram-positive commensals in the gut and may point to a lipid-mediated axis of host–pathogen interaction^48^.

Among gene products specifically upregulated during superinfection, alcohol dehydrogenases emerged as essential. *AdhE*, and to a lesser extent *adhA*, were robustly and consistently induced across time points and serotypes, and proved indispensable for pneumococcal fitness specifically in the IAV-modified lung. Importantly, deletion of either gene alone had no measurable impact, whereas combined deletion resulted in a striking defect during superinfection but not during primary pneumococcal pneumonia. This complete functional redundancy, coupled with context-specific essentiality, distinguishes AdhE/A from classical virulence determinants or housekeeping metabolic enzymes that are broadly required *in vivo*^21^.

On the host side our data identify two major, synergistic driving forces that converge on this metabolic dependency: altered carbon source availability and reduced oxygen tension. Untargeted metabolomic profiling revealed an accumulation of alternative carbohydrates, most prominently maltose and mannose, in bronchoalveolar lavage cells following IAV infection. These findings expand on prior work showing glucose leakage into the alveolar space during influenza^39^, and suggest that the post-viral lung is characterized not simply by nutrient abundance, but by qualitative changes in the carbon landscape. *S. pneumoniae* is metabolically versatile and capable of co-utilizing multiple carbohydrates^49^ even in the presence of glucose^50^, particularly under conditions where classical carbon catabolite repression is attenuated^51^. Consistent with this, we observed robust *in vivo* upregulation of maltodextrin transport and utilization genes, and we demonstrate that maltose, maltotriose, glycogen, and other non-glucose sugars are sufficient to induce *adhE* expression *in vitro*.

The origin of these alternative carbohydrates remains speculative but likely reflects virus- and inflammation-induced tissue remodeling. Glycogen is abundant in airway epithelial cells and immune cells and accumulates in the lung during tissue damage^52^ . Pneumococcal enzymes such as SpuA can break down host glycogen to malto-oligosaccharides which are then imported and metabolized through dedicated systems^40,53^. The coordinated induction of maltodextrin uptake systems and alcohol dehydrogenases suggests a model in which pneumococcus responds to and exploits complex host-derived carbohydrates that become accessible during viral lung injury, thereby reshaping its central metabolism.

In parallel, our data support a critical role for hypoxia in driving pneumococcal metabolic reprogramming during superinfection. Host transcriptomic analyses revealed induction of hypoxia-responsive genes following IAV infection, consistent with prior reports of localized hypoxia in infected lungs^43,54,55^. Bacteria respond by repression of oxygen-dependent pathways such as SpxB and enhanced reliance on fermentative metabolism. Under these conditions, regeneration of NAD⁺ becomes a limiting factor for sustaining glycolytic flux. Alcohol dehydrogenases provide a solution to this redox constraint by enabling mixed-acid fermentation and NADH re-oxidation when lactate dehydrogenase and oxygen-dependent systems are insufficient. Our *in vitro* experiments confirm that anaerobiosis and alternative carbon sources act synergistically to induce adh expression, closely recapitulating the *in vivo* transcriptional signature observed during superinfection. Of note, anaerobic culture of pneumococcus *in vitro* was previously shown to enhance its virulence and dissemination *in vivo*^56^, possibly due to preemptive establishment of the necessary transcriptional program prior to transfer in the host. Interestingly, interference with Hif1a alleviates pathology during superinfection, indicating that hypoxia likely contributes to disease not only by shaping bacterial metabolism but also by engaging host-driven pathogenic programs^57^. *adhE* may consequently play a relevant role in infection of patients with chronic respiratory conditions such as asthma or COPD, which are associated with hypoxia and heightened sensitivity to pneumococcal infection^58,59^.

From a therapeutic perspective, this convergence reveals a context-specific metabolic vulnerability. Although targeting bacterial carbohydrate utilization has been proposed as an antimicrobial strategy, for example through sugar analogs^60^ the marked metabolic flexibility of *S. pneumoniae* is likely to limit the efficacy of approaches focused on individual nutrient pathways. Instead, our data identify alcohol dehydrogenase–mediated NAD⁺ regeneration as a downstream metabolic bottleneck that arises specifically in the virus-modified pulmonary environment and is shared across pneumococcal strains.

Consistent with this, inhibition of alcohol dehydrogenases using the FDA-approved drug fomepizole selectively restricted pneumococcal growth and dissemination during superinfection, while having no effect during primary pneumonia. This specificity highlights the value of targeting adaptive, context–dependent bacterial processes rather than core metabolic functions and suggests that virus-induced remodeling of the lung creates transient metabolic liabilities that can be therapeutically exploited. Importantly, fomepizole has previously been evaluated in preclinical models of pneumococcal infection, where it showed little effect on bacterial expansion when administered alone, consistent with our data, but synergized with antibiotic treatment by enhancing bacterial redox stress^26^. Thus, in the context of antibiotic treatment, fomepizole could potentially be repurposed in pneumococcal infection, independently of prior viral exposure.

More broadly, given the prevalence of secondary bacterial infections following diverse respiratory viral infections, including SARS-CoV-2 and RSV, similar context-dependent metabolic dependencies may extend across bacterial pathogens and viral settings^61–63^.

In summary, by disentangling adaptive bacterial responses from generic virulence programs, we establish alcohol dehydrogenase–mediated redox metabolism as a central determinant of superinfection and reveal conserved, temporally restricted vulnerabilities that may be leveraged to limit secondary bacterial disease.

## Methods

### Bacterial strains and culture conditions

*S. pneumoniae* (D39L) was kindly provided by Prof. Marien De Jonge and *S. pneumoniae* (ATCC-6303) was purchased from LGC (Germany). Both were cultivated in Trypticase soy broth (TSB) and on Trypticase soy agar plates with 5% sheep blood (Thermo Scientific) at 37°C in 5 % CO2. When required, media where supplemented with kanamycin (500 µg/mL), spectinomycin (100 µg/mL) and colistin (2 µg/mL). For *in vitro* stimulation with different sugars, *S. pneumoniae* was grown in a chemically defined medium (CDM) prepared as previously described^64^. Culture in anaerobic conditions were performed in a vinyl anaerobic chamber (Coy Laboratory Products, gas composition: 85% N₂, 10% CO₂ and 5% H₂). All media used under anaerobic conditions were pre-reduced before use.

### Mouse strains and husbandry

C57BL/6J mice (female, 7–10 weeks of age) were purchased from Charles River Laboratories (France). Mice were housed under SPF/BSL2 conditions with a 12-h light/dark cycle and fed ad libitum. All animal experiments were conducted in accordance with legal and ethical regulations. Experimental procedures were approved by the Department of Animal Experimentation of the Geneva canton under license numbers GE168A-GE168B and GE513.

### Animal infection

For all infections, mice were anaesthetized by intraperitoneal (i.p.) injection of 200 µl sterile PBS containing ketamine/xylazine (100 mg kg⁻¹ and 10 mg kg⁻¹, respectively). At day 0, mice were inoculated intranasally (i.n.) with 40 µl of either PBS or influenza A/Netherlands/602/2009 (Neth/602) at a dose of 10 PFU. Seven days later, mice were infected i.n. with 40 µl of *S. pneumoniae*. IAV-primed mice were challenged with a low bacterial dose of 1 × 10³ CFU for ATCC-6303 or 1 × 10⁴ CFU for the D39L strain, whereas PBS-primed mice received either the same low dose (Spn Low) or a higher dose of 1 × 10⁶ CFU (ATCC-6303) or 1× 10⁸ CFU (D39L) (Spn). Mice in the IAV-only group were inoculated with IAV at day 0 and received PBS at day 7. Animals were euthanized 24 h after *S. pneumoniae* infection by i.p. administration of 150 mg kg⁻¹ pentobarbital. Lungs were collected and either preserved in RNAprotect Tissue Reagent (Qiagen) for RNA extraction or homogenized in PBS containing 0.01% Triton X-100 for CFU enumeration.

To study superinfection over time, mice were inoculated with PBS or IAV at day 0 and subsequently infected with *S. pneumoniae* at day 7, 14, or 28, and sacrificed 24 h later (corresponding to weeks 1, 2 and 4, respectively). To assess early bacterial adaptation, IAV-primed mice were infected at day 7 with high-dose *S. pneumoniae* and euthanized 2 h post-infection.

For the *in vivo* time-course experiment, mice primed with IAV at day 0 were subsequently infected with 1 × 10⁴ CFU of *S. pneumoniae* (SI) at day 7 and were sacrificed at 2, 5, 8, 12, 15, 18 or 21 hours post bacterial infection. PBS-primed mice at day 0 were subsequently infected with 1 × 10⁴ CFU (Spn low) or 1 × 10^8^ CFU of *S. pneumoniae* (Spn) and sacrificed at 2 or 5 hours post bacterial infection.

### *In vivo* competitive index

One week after IAV infection or PBS priming, mice were infected intranasally with a 1:1 mixture of wild-type (WT) and knockout (KO) *S. pneumoniae*. As described above, PBS-primed mice received 1 × 10⁸ CFU of the WT/KO mixture, whereas IAV-primed mice received 1 × 10⁴ CFU. 24 h after bacterial infection, mice were euthanized and lungs were collected in PBS containing 0.01% Triton X-100. Lungs were homogenized, serially diluted and plated on Columbia agar supplemented with 5% sheep blood (COS) containing either colistin to suppress resident lung microbiota or a selective antibiotic for mutant recovery. The competitive index (CI) was calculated as the ratio of KO to WT bacteria recovered from the lungs. In rare cases, KO bacteria outcompeted the WT strain such that no WT colonies were recovered from lung homogenates. When this occurred, a maximal CI value of 20 was assigned for analysis. This value was determined by representative cases in which the KO/WT ratio was verified using colony PCR and strain-specific qPCR.

### Alcohol dehydrogenase inhibition *in vivo*

Using the week 1 infection protocol, mice were injected intraperitoneally with either PBS (control) or 4-methylpyrazole (Fomepizole; ThermoFisher Scientific) at 50 mg kg^-1^. Fomepizole was administered at the time of *S. pneumoniae* infection, mixed with the anesthetic solution, and again 12 h post-infection. The dosing regimen was based on previously reported use of Fomepizole in the context of *S. pneumoniae* infection^26^. Mouse lungs and spleens were harvested to determine bacterial CFU.

### Blocking antibody injection *in vivo*

Using the week 1 infection protocol, mice were injected intraperitoneally with either an isotype control antibody or IL-21R–blocking or IFN-g–blocking antibodies (clones 4A9 and XMG1.2, respectively; Bio X Cell). Antibodies were administered at 200 µg in 200 µl of PBS per dose on days 5, 6 and 7 post-IAV infection. The final injection on day 7 was performed at least 4 h before *S. pneumoniae* infection to avoid interference with the anesthetic agents.

### Lung RNA extraction

Lungs preserved in RNAprotect Tissue Reagent (Qiagen) were transferred into 1 mL of TRIzol (Invitrogen), bead homogenized to break down tissue (Speed 6, 30 sec). The lysate was transferred in Bead Tubes for PureLink Microbiome DNA Purification Kit (Invitrogen) and bead homogenized to break down host and bacteria (speed 4.30 for 1 min, with two 15-s breaks). Then total RNA was extracted using TRIzol following manufacturer’s instruction. Briefly, 0.2 mL of chloroform was added, samples were incubated 10 min at room temperature and centrifuged at 12,000 × g at 4°C for 15 min. The aqueous phase was transferred into a new tube, mixed with 0.5 mL of isopropanol and incubated 10 min at 4°C. Samples were centrifuged 10 min at 4°C and RNA pellets were washed twice with 75% ethanol and centrifuged 5 min at 7,500 × g at 4°C. Pellets were air-dried and resuspended in 200 µl of RNase-free water.

### qRT-PCR on lung samples

Bulk lung RNA was treated with DNase using the DNA-free Kit (Invitrogen), and RNA concentrations were determined with the Qubit RNA High Sensitivity Assay (Thermo Fisher Scientific). cDNA synthesis was performed with random primers and SuperScript II (Invitrogen) following the manufacturer’s instructions. qPCR reactions were assembled in Hard-Shell 96-well PCR plates (Bio-Rad), sealed with Microseal B film (Bio-Rad) and run on CFX96 Real-Time PCR Detection System (Bio-Rad) using PowerUp SYBR Green Master Mix (Applied Biosystems). Primers are listed in Supplementary Table 2. qPCR data were normalized to the housekeeping gene gyrB and to the mean of the control samples, and are expressed as log₂ fold change.

### *S. pneumoniae* CFU estimation from lung by qRT-PCR

Estimated CFU were used to determine the relative abundance of *S. pneumoniae* in the different samples. To obtain CFU equivalents from lung samples, Ct values obtained for the bacterial housekeeping gene *gyrB* were compared to a standard curve generated from *in vitro* cultures with known CFU. For each point on the standard curve, the corresponding CFU value was calculated based on the amount of cDNA used per reaction and the total RNA yield obtained from the *in vitro* culture. This standard curve was then used to convert *gyrB* Ct values from lung-derived cDNA into estimated CFU equivalents for each sample.

### Dual RNA sequencing and analysis

RNA quality was assessed using a Bioanalyzer (Agilent Technologies). Libraries were prepared with the Ribo-Zero Plus rRNA Depletion Kit (Illumina) and sequenced on an Illumina NovaSeq 6000 platform, generating 100 bp single-end reads. Sequencing depth ranged from approximately 100– 300 million reads per sample for bacteria-containing samples and ∼30 million reads for bacteria-negative samples. RNAseq reads were mapped with STAR v2.7.4 independently on each reference genome; mouse genome (GRCm39 from Genecode vM27), Spn D39L (assembled and annotated with PGAP version 2023-10-03.build7061), Spn ATCC6303 (genome sequence downloaded from ATCC, and annotated with PGAP version 2023-10-03.build7061), and IAV genome (8 segments sequences and annotations downloaded from NCBI: CY176942.1, CY176943.1, CY176944.1, CY176945.1, CY176946.1, CY176947.1, CY176948.1, CY176949.1). The computations were performed on the HPC cluster of the University of Geneva. All read quantification were done with R programming language using the *GenomicAlignments::summarizeOverlaps* method *(inter.feature = FALSE),* tuned to count antisense reads for each gene. Samples with insufficient *S. pneumoniae* read coverage were excluded from downstream transcriptomic analyses. For the mouse genome, exons split by genes were given to quantify antisense exonic reads only. Orthologous genes between Spn strains were identified by tblastx pairwise alignment of genes sequences retaining only reciprocal best hits. KEGG annotations of our genes were obtained by submitting gene sequences to BlastKOALA service.

### Genomic DNA sequencing

Genomic DNA was extracted using the Wizard HMW DNA Extraction Kit (Promega). Long-read sequencing libraries were prepared with the Native Barcoding Kit 24 V14 (Oxford Nanopore Technologies) and sequenced on a MinION Mk1C using an R10.4.1 flow cell (Oxford Nanopore Technologies). Raw reads were basecalled using the Guppy basecaller (v.6.4.2+97a7f06) with the super-accuracy model (dna_r10.4.1_e8.2_400bps_sup). De novo genome assemblies were generated with Flye (v2.9.1), using the longest 1% of reads to construct the initial assembly backbone. Assemblies were then polished with Medaka (v1.7.0) to generate the final consensus sequences.

### *S. pneumoniae* knockout mutant construction

Single-gene deletion mutants were generated by replacing the target locus with a kanamycin-resistance cassette, and a double-knockout strain was obtained by introducing a spectinomycin-resistance cassette into the *adhA* knockout background. Kanamycin and spectinomycin cassettes were amplified using Q5 High-Fidelity DNA Polymerase (New England Biolabs) from the Janus cassette ^65^ kindly provided by Prof. Lucy Hathaway and from the pASR102 ^66^, respectively. Each antibiotic cassette was cloned between ∼1 kb upstream and downstream homology arms flanking the target gene and assembled using HiFi DNA Assembly (New England Biolabs). Primers used for plasmid construction are listed in Supplementary Table 2.

Linear DNA constructs containing the antibiotic-resistance cassette and both homology arms were amplified with Q5 and purified with the QIAquick PCR Purification Kit (Qiagen) prior to transformation into *S. pneumoniae*. Transformation was performed as previously described^67^. Briefly, competent cells were prepared by inoculating a single colony into brain heart infusion (BHI) broth supplemented with 5% FCS and growing the culture overnight at 37 °C. The culture was diluted into pre-warmed BHI–FCS and grown to an OD_600nm_ of ∼0.15, then diluted 1:20 into competence medium (TSB (pH 8.0), 10% glycerol, 0.16% bovine serum albumin, 0.01% CaCl2) pre-warmed to 30 °C. 1 ml of bacterial aliquots were incubated for 15 min at 30 °C, followed by addition of 100 ng CSP-1 and a further 15 min incubation before addition of 500 ng of linear DNA. Cultures were incubated for 40 min at 30 °C and then for 90 min at 37 °C to allow expression of antibiotic resistance. Cells were plated on Columbia sheep blood agar (Thermo Fisher Scientific) containing the appropriate antibiotic and incubated overnight at 37 °C in 5% CO_2_. Transformant clones were verified by PCR screening, and total RNA was extracted from confirmed mutants to validate loss of gene expression by qRT–PCR.

### Sugar stimulation of bacterial culture in aerobic and anaerobic conditions

*S. pneumoniae* was grown in TSB up to an OD_600nm_ of 0.4 and washed in CDM lacking carbon sources (without pyruvate and glucose). Bacteria were incubated for 30 min in carbon source-deprived CDM, then 1% (w/v) sugars were added to the bacterial culture and incubated for 30 min. To test the impact of oxygen on *S. pneumoniae* gene expression, cultures under aerobic and anaerobic conditions were performed simultaneously.

### Hot phenol RNA extraction and qRT-PCR of bacterial RNA

Bacterial cultures were stopped by adding STOPmix solution (20% of the sample volume; 95% ethanol, 5% phenol saturated with 0.1 M citrate buffer, pH 4.3), mixed by inversion and snap-frozen in liquid nitrogen. RNA was extracted using the hot phenol RNA extraction protocol as previously described ^48^. Reverse transcription was performed using the QuantiTect RT Kit (Qiagen) according to the manufacturer’s instructions. qPCR reactions were assembled in MicroAmp 96- or 384-well optical plates (Applied Biosystems), sealed with MicroAmp Optical Adhesive Film and run on a QuantStudio 6 Pro system (Applied Biosystems). Reactions were prepared using PowerUp SYBR Green Master Mix (Applied Biosystems) following the manufacturer’s guidelines. Primers are listed in Supplementary Table 2. qPCR data were normalized to the housekeeping gene *gyrB* and to the mean of control samples, and are expressed as log₂ fold change.

### Bronchoalveolar lavage fluid collection

Bronchoalveolar lavage fluid (BALF) was collected from mice at the time of sacrifice. Lungs were lavaged by gently instilling sterile, ice-cold phosphate-buffered saline (PBS) via a tracheal cannula. A volume of 1 mL PBS was instilled and slowly withdrawn; this procedure was repeated once, and recovered lavage fluids were pooled for each mouse. BALF was centrifuged at 400 × g for 5 min at 4 °C to pellet cells. The resulting supernatant was subsequently centrifuged at 9,600 × g for 5 min at 4 °C to remove residual debris and potential commensal bacteria. Both the cell pellet and BALF supernatant were snap-frozen in liquid nitrogen and stored until metabolite analysis.

### Metabolite Profiling by Gas Chromatography-Mass Spectrometry (GC-MS)

To extract and separate metabolites from cell pellets, 50 µl chloroform were added to cell pellets, followed by 200 µl of a methanol/water mix (3:1, all solvents used were of mass spectrometry grade). All samples were vigorously mixed at 4 °C for 30 min. Subsequently, samples were centrifuged (20,000 × g, 6 min, 4 °C) and the supernatant was transferred to microfuge tubes containing 100 µl water, resulting in a chloroform/methanol/water ratio of 1:3:3 (biphasic). Samples were vigorously mixed again at 4 °C for 10 min before centrifuging samples (20,000 × g, 6 min, 4 °C). The upper polar phase was transferred to new microfuge tubes and dried sequentially inside mass spectrometry vial inserts using a centrifugal evaporator (Savant Speed Vac Concentrator). The extract was further dried and concentrated through addition of decreasing volumes of methanol. To the dried, concentrated metabolite extracts, 30 µl pyridine (Sigma-Aldrich, 270970) were added, containing methoxyamine hydrochloride (20 mg/ml, Sigma-Aldrich, 226904). Methoximation was allowed to proceed overnight at room temperature, while rocking the samples gently. The next day, 30 µl of N,O-Bis(trimethylsilyl)trifluoroacetamide with 1% trimethylchlorosilane (BSTFA TMCS, TMS, Supelco, B-023) were added for silylation. Samples were vortexed and analyzed at least 30 min after addition of TMS, using an 8890 GC System (Agilent) equipped with a DB5 capillary column (JCW Scientific, 30 m, 250 μm inner diameter, 0.25-μm film thickness), with a 10-m inert Duraguard, connected to a 5977B GC/MSD operated in electron impact (EI, 70 eV) mode linked to a 7693A autosampler (Agilent). The GC–MS settings were as follows: Inlet temperature: 270 °C, MS transfer line temperature: 280°C, MS source temperature: 230°C and MS quadrupole temperature: 150 °C. The oven temperature gradient during the sample run was as follows: 80 °C (2 min); 80 °C to 140 °C at 30 °C/min; 140 °C to 250 °C at 5 °C/min; 250 °C to 310 °C at 15 °C/min; 310 °C for 2 min. All samples were analyzed twice in a randomized sequence following injection of 3 μl (splitless) with a solvent delay of 7 min, once in scan mode (*m/z* 80–800), and once in selected ion monitoring (SIM) mode. In SIM mode, ions of the internal standard (^13^C3/^15^N β-alanine, *m/z* 176, 250 and 294) as well as prominent ions of relevant disaccharides (maltose, sucrose and lactose, *m/z* 361, 217, 204) were monitored. Metabolites were identified based on the retention time and presence of three identifier ions, determined by the analysis of authentic standards. The identifier ions of the internal standard and disaccharides were as described above. Metabolite abundances were determined across all samples using MassHunter Quantitative Analysis Software 12.0 (Agilent), quantifying a suitable ion at the determined peak apex with a left and right retention time delta of 0.06 min or 0.1 min for very abundant metabolites (e.g., glucose). The ions *m/z* 250 and *m/z* 361 were selected as quantifier ions for the internal standard and disaccharides, respectively. The identifier and quantifier ions of other relevant metabolites were as follows, with the quantifier ion listed first: arabinose (*m/z* 307, 217, 103), galactose (*m/z* 319, 205, 147), glucose (*m/z* 319, 205, 160), mannose (*m/z* 319, 205, 147). Metabolite abundances were normalized to the total ion chromatogram (TIC) intensity of the respective sample, and normalized mock metabolite abundances were set to a relative abundance of 1.

### Flow cytometry

Following sacrifice mice were injected intravenously with anti-CD45 antibody to label circulating immune cells. Perfused murine lungs were mechanically homogenized with gentleMACS™ Octo Dissociator (Miltenyi Biotec) in 5 ml PBS, 5% FBS, 2 mg mL^-1^ collagenase IV (Worthington) and 0.1 mg/ml DNase. The homogenized tissue was incubated shaking for 40 min at 37 ℃, then passed through a 70 µm strainer and red blood cells were lysed. Immune cells were enriched using 40% Percoll (GE) and washed with PBS. The recovered cells were incubated with TruStain FcX™ (anti-mouse CD16/32) Antibody (Biolegend) and stained with the antibodies listed in Extended Data table 3. All analyses were performed with BD Fortessa and FlowJo software.

### Statistical analysis

All statistical analyses were performed using GraphPad Prism (version 10). Details of each experiment, including the statistical tests used, the definition of *n*, and measures of central tendency and dispersion, are provided in the corresponding figure legends. Comparisons between two groups were performed using two-tailed unpaired Student’s t-tests. Comparisons among more than two groups were performed using one-way analysis of variance (ANOVA) followed by appropriate post hoc multiple-comparison tests (Tukey’s, Šidák’s, or Dunnett’s). No formal tests of normality were performed due to limited sample sizes. A *P* value < 0.05 was considered statistically significant. Levels of significance are indicated in the figures and legends as follows: ns, not significant; *P* < 0.05; P < 0.01; *P* < 0.001; and P < 0.0001.

## Acknowledgments

We thank all members of our laboratories. We thank the iGE3 Genomic Platform, the animal facility and animal caretaker of the University of Geneva. We thank Prof. Lucy Hathaway for kindly for providing the plasmid containing the kanamycin cassette and *S. pneumoniae* transformation guidance. Prof. Marien de Jonge for providing the D39L strain.

This project was supported by the Fondation privée des HUG (CONFIRM#RC07-07) and by the Novartis foundation (NOVARTIS #24B142). S.B. was further supported by an Eccellenza Professorial Fellowship (PCEFP3_187018) and a Starting Grant (TMSGI3_211235) from the Swiss National Science Foundation.

## Author contributions

M.L. designed and performed most of the experiments, with assistance from A.C., G.K., and F.S., and analyzed data; N.B. performed all FACS analyses; J.P. conducted the bioinformatic analyses; R.S. sequenced and analyzed *S. pneumoniae* genome; J.K. performed metabolomic analyses with assistance from O.V.R.; J.W.V. provided critical reagents and intellectual input; M.C. performed metabolomic analyses under the supervision of M.S.G. and M.S.G. provided intellectual input. M.L., M.S. and S.B. wrote the paper, with inputs from all authors; M.S. and S.B. designed and supervised the study.

