## Extended data figures for "Early metabolic reprogramming licenses *Streptococcus pneumoniae* for Influenza-driven superinfection"

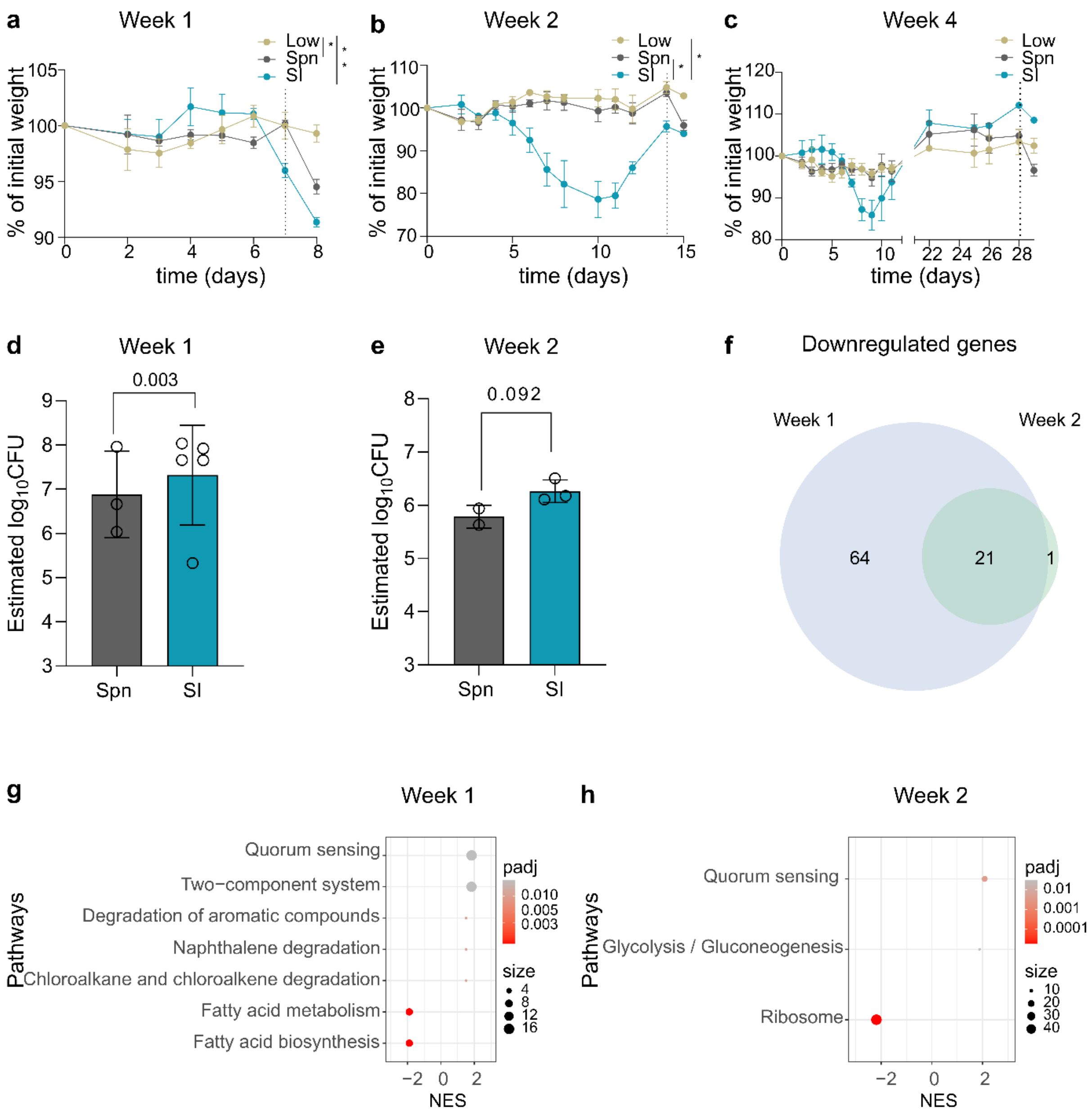

a
Week 1
Low
Spn
SI
% of initial weight
time (days)
b
Week 2
c
Week 4
d
Week 1
0.003
Estimated log10CFU
Spn
SI
e
Week 2
0.092
f
Downregulated genes
Week 1
Week 2
64
21
1
g
Week 1
Pathways
Quorum sensing
Two−component system
Degradation of aromatic compounds
Naphthalene degradation
Chloroalkane and chloroalkene degradation
Fatty acid metabolism
Fatty acid biosynthesis
padj
size
NES
h
Week 2
Quorum sensing
Glycolysis / Gluconeogenesis
Ribosome

**Extended data figure 1: *S. pneumoniae* adaptation to IAV modified lung environment over time**

**a-c**, Percentage of initial body weight of mice infected with ~1 × $10^8$ CFU *S. pneumoniae* (Spn) or ~1 × $10^4$ CFU (Spn Low) and superinfected (SI) at **(a)** one week, **(b)** two weeks or **(c)** four weeks following IAV infection or PBS priming. The dotted line indicates the time of *S. pneumoniae* infection. Statistical significance was determined at **(a)** day 7 and **(b)** day 10. Data are means ± s.e.m. (n = 2–3 per group; one representative of at least two experiments is shown). **d,e,** Estimation of bacterial titer at **(d)** one week and **(e)** two weeks following IAV infection, based on expression of the housekeeping gene *gyrB,* as determined by RT-qPCR. Dots indicate individual mice (n = 2-5 per group; geometric means ± geometric s.d. are shown). **f**, Venn diagram showing *S. pneumoniae* genes downregulated one week and two weeks following IAV infection. **g,h**, Top activated and suppressed *S. pneumoniae* gene sets at **(g)** one week and **(h)** two weeks following IAV infection (KEGG pathways). Statistical significance was determined by two-way ANOVA **(a-c)** and two-tailed unpaired *t*-tests **(d,e)**.

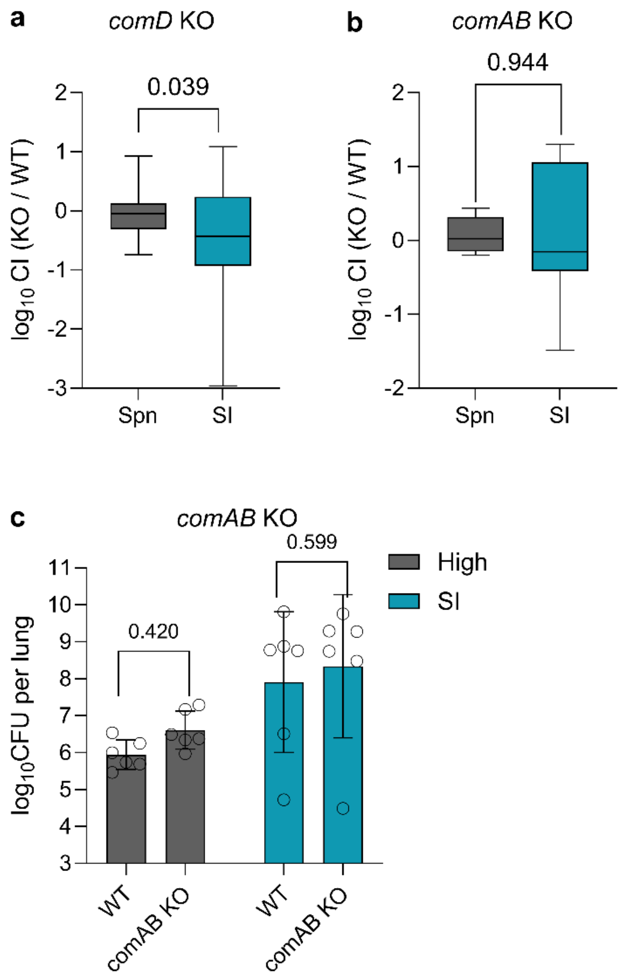

a
comD KO
0.039
log10 CI (KO / WT)
2
1
0
-1
-2
-3
Spn
SI
b
comAB KO
0.944
log10 CI (KO / WT)
2
1
0
-1
-2
Spn
SI
c
comAB KO
0.599
0.420
High
SI
log10CFU per lung
11
10
9
8
7
6
5
4
3
WT
comAB KO
WT
comAB KO

**Extended data figure 2 : Isogenic knock-out mutants of the *S. pneumoniae* competence system show no fitness defect during superinfection**

**a,b**, Competitive infection with WT and isogenic deletion mutant **(a)** *comD* KO and **(b)** *comAB* KO. Mice primed with PBS or IAV one week prior were infected with a 1:1 ratio of KO to WT bacteria, and sacrificed 24 h later to determine relative burden. (**a**, n = 6-9 per group; one experiment; **b,** n= 23-28 per group, data are pooled from at least 2 independent experiments; box-and-whisker plots show the median (centre line), interquartile range (box), and minimum and maximum values (whiskers)). **c**, Bacterial burdens 24 h post individual infection with WT or *comAB* KO strains in mice primed with PBS (Spn) or IAV (SI) seven days prior. Dots represent individual mice (n= 5-6 per group, one experiment; geometric means ± geometric s.d. are shown). Statistical significance was determined by two-tailed unpaired *t*-tests **(a,b)** and two-way ANOVA **(c)**.

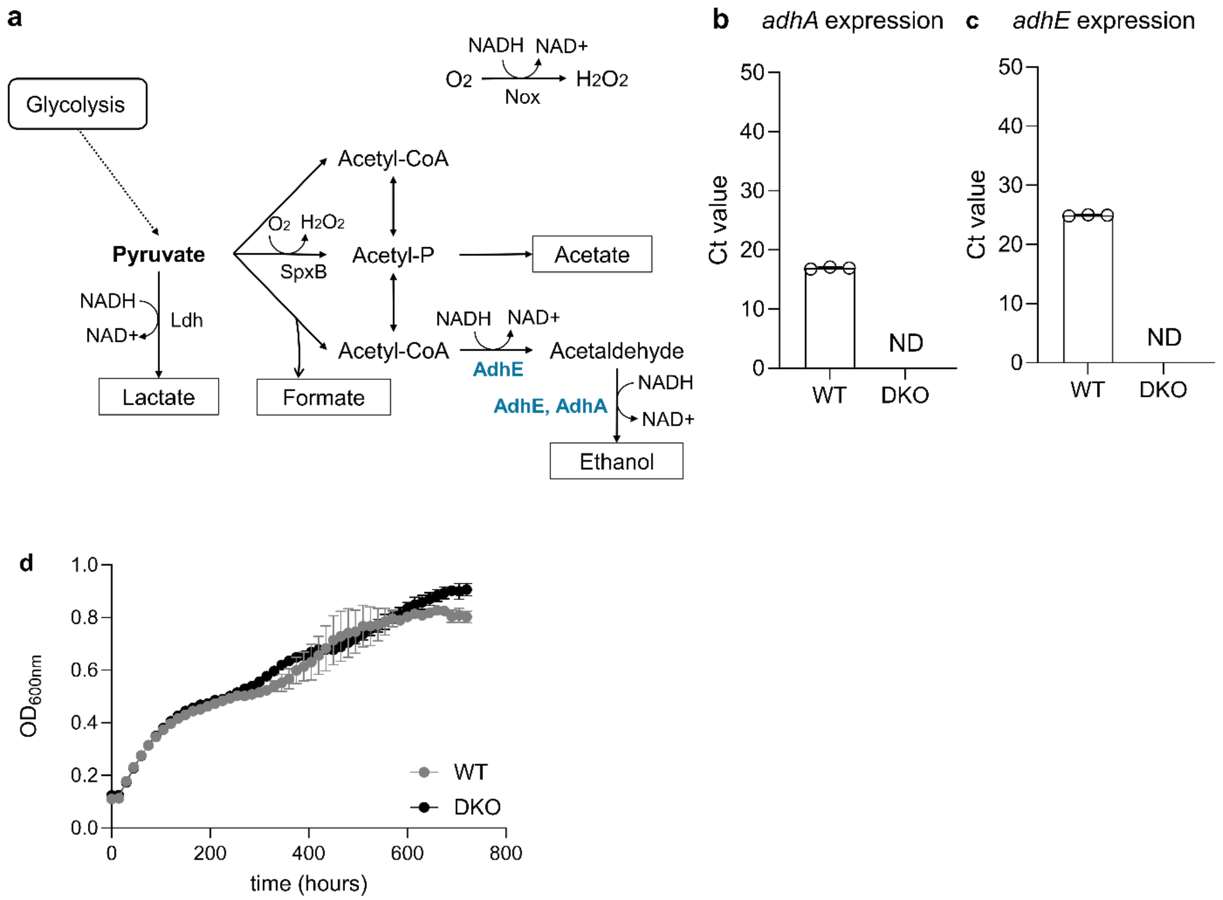
a
Glycolysis
Pyruvate
NADH
NAD+
Ldh
Lactate
O2 H2O2
SpxB
Formate
Acetyl-CoA
Acetyl-P
Acetyl-CoA
Acetate
NADH NAD+
O2
H2O2
Nox
NADH NAD+
AdhE
Acetaldehyde
NADH
AdhE, AdhA
NAD+
Ethanol
b
adhA expression
Ct value
WT
DKO
ND
c
adhE expression
Ct value
WT
DKO
ND
d
OD600nm
time (hours)
WT
DKO

**Extended data figure 3: *adhE/adhA* DKO construct**

**a**, Schematic of the pyruvate metabolism and NAD+ regeneration in *S. pneumoniae*. **b,c**, RT-qPCR analysis of **(b)** *adhA* and **(c)** *adhE* expression, shown as Ct values in WT and *adhE/adhA* DKO. Dots represent individual bacterial cultures (n = 3 per group, one representative of at least two experiments is shown; means ± s.d. are shown). **d**, *in vitro* growth curves of WT and DKO in TSB medium (n = 3 per group; one representative of at least two experiments is shown; means ± s.e.m. are shown).

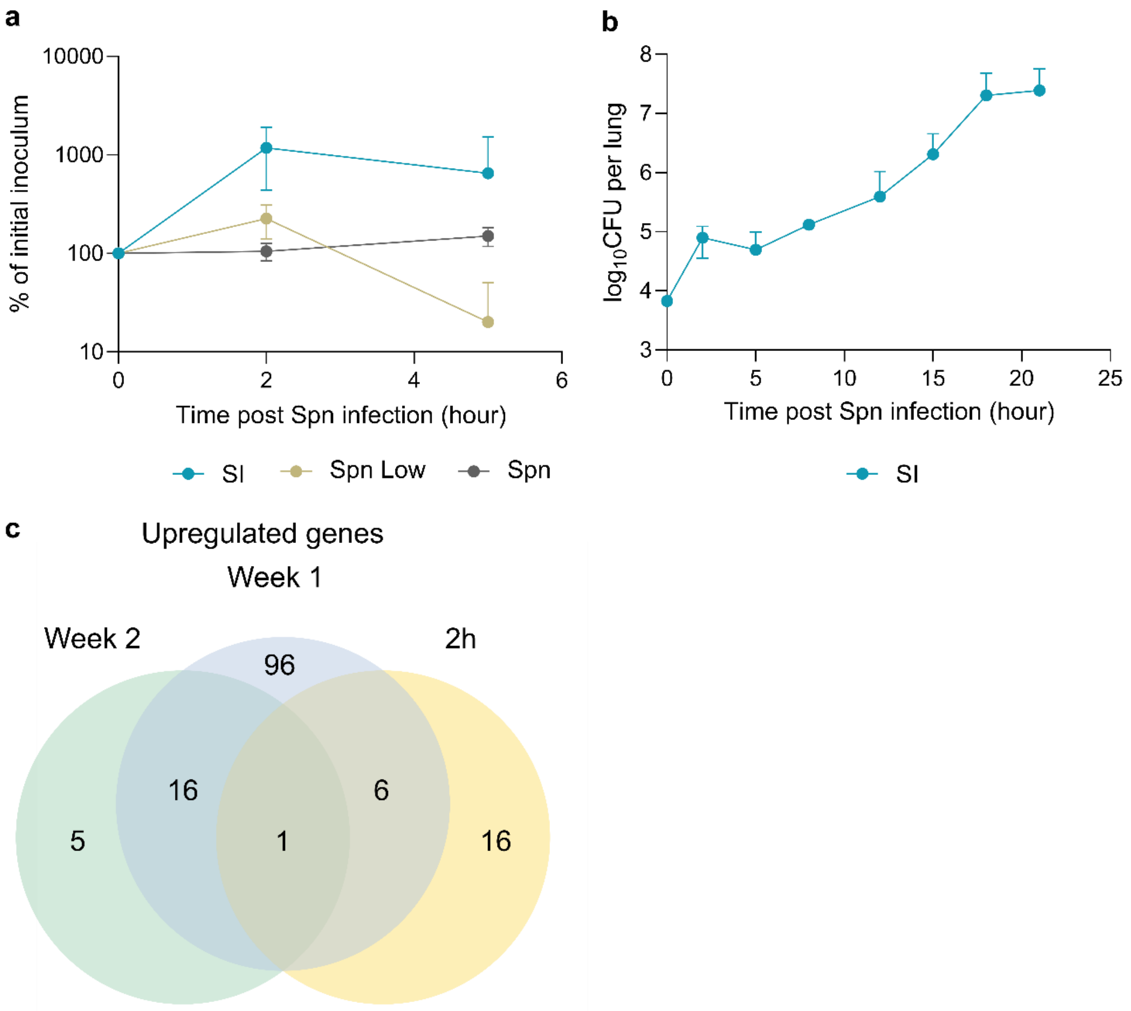
a
10000
1000
100
10
% of initial inoculum
0
2
4
6
Time post Spn infection (hour)
SI
Spn Low
Spn
b
8
7
6
5
4
3
log10CFU per lung
0
5
10
15
20
25
Time post Spn infection (hour)
SI
c
Upregulated genes
Week 1
Week 2
2h
96
16
6
5
1
16

**Extended data figure 4: *S. pneumoniae* adaptation to IAV modified lung environment at early time point (2h post-bacterial infection)**

**a**, Percentage of initial inoculum retrieved in mouse lungs following Spn Low, Spn and SI infection (n = 2-4 per group and time point; data pooled from 2 independent experiments; means ± s.d. are shown). **b,** Bacterial titers in SI mice over time (n = 2-5 per time point; data pooled from two independent experiments; means ± s.d. are shown). **c**, Venn diagram of gene upregulated during superinfection at one week and two weeks post IAV infection and at early time point (2 h post *S. pneumoniae* infection).

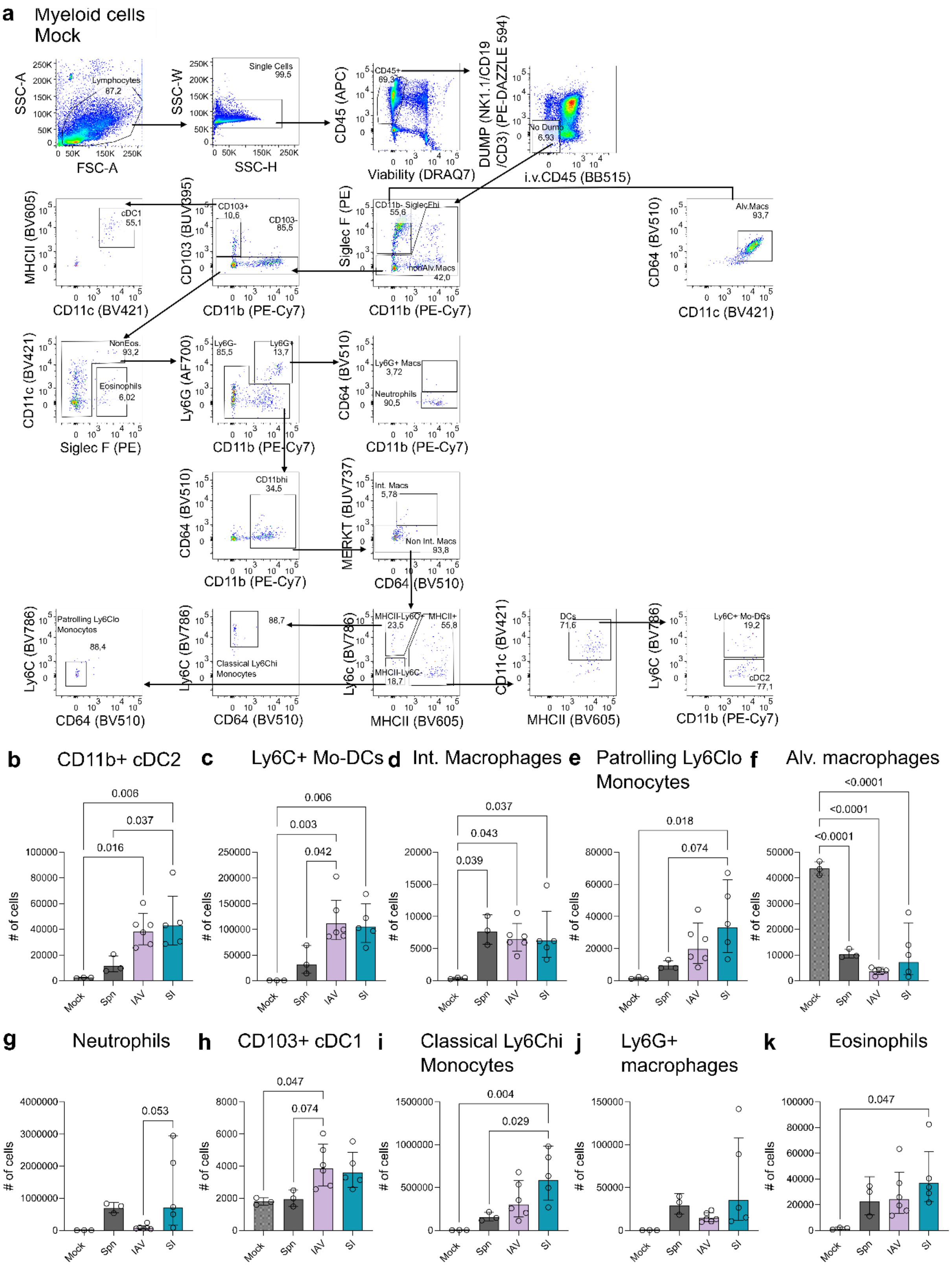
a
Myeloid cells
Mock
SSC-A
FSC-A
Lymphocytes 87,2
SSC-W
SSC-H
Single Cells 99,5
CD45 (APC)
Viability (DRAQ7)
CD45+ 69,3
DUMP (NK1.1/CD19 /CD3) (PE-DAZZLE 594)
i.v.CD45 (BB515)
No Dump 6,93
MHCII (BV605)
CD11c (BV421)
cDC1 55,1
CD103 (BUV395)
CD11b (PE-Cy7)
CD103+ 10,6
CD103- 85,5
Siglec F (PE)
CD11b- SiglecFhi 55,6
nonAlv.Macs 42,0
CD64 (BV510)
Alv.Macs 93,7
CD11c (BV421)
Siglec F (PE)
NonEos. 93,2
Eosinophils 6,02
Ly6G (AF700)
Ly6G- 85,5
Ly6G+ 13,7
Ly6G+ Macs 3,72
Neutrophils 90,5
CD11bhi 34,5
MERKT (BUV737)
CD64 (BV510)
Int. Macs 5,78
Non Int. Macs 93,8
Ly6C (BV786)
Patrolling Ly6Clo Monocytes 88,4
88,7
Classical Ly6Chi Monocytes
Ly6c (BV786)
MHCII (BV605)
MHCII-Ly6C+ 23,5
MHCII+ 55,8
MHCII-Ly6C- 18,7
DCs 71,6
Ly6C+ Mo-DCs 19,2
cDC2 77,1
b CD11b+ cDC2
c Ly6C+ Mo-DCs
d Int. Macrophages
e Patrolling Ly6Clo Monocytes
f Alv. macrophages
g Neutrophils
h CD103+ cDC1
i Classical Ly6Chi Monocytes
j Ly6G+ macrophages
k Eosinophils
### of cells
Mock
Spn
IAV
SI
0.006
0.037
0.016
0.003
0.042
0.037
0.043
0.039
0.018
0.074
<0.0001
<0.0001
<0.0001
0.053
0.047
0.074
0.004
0.029
0.047

**Extended data figure 5: Influenza infection induces myeloid cell infiltration in lung at day 8 post infection**

**a,** Gating strategy of myeloid cells. **b-k**, Number of cells in Mock, IAV, Spn and SI infected mice. **(b)** CD11b+ cDC2, **(c)** Ly66C+ monocyte derived dendritic cells, **(d)** interstitial macrophages, **(e)** patrolling Ly6Clo monocytes, **(f)** alveolar macrophages, **(g)** neutrophils, **(h)** CD103+ cells cDC1, **(i)** classical Ly6C high monocytes, **(j)** Ly6G+ macrophages, **(k)** eosinophils (n = 3-6 per group). Data are shown as geometric means ± geometric s.d. Statistical significance was determined by ordinary one-way ANOVA.

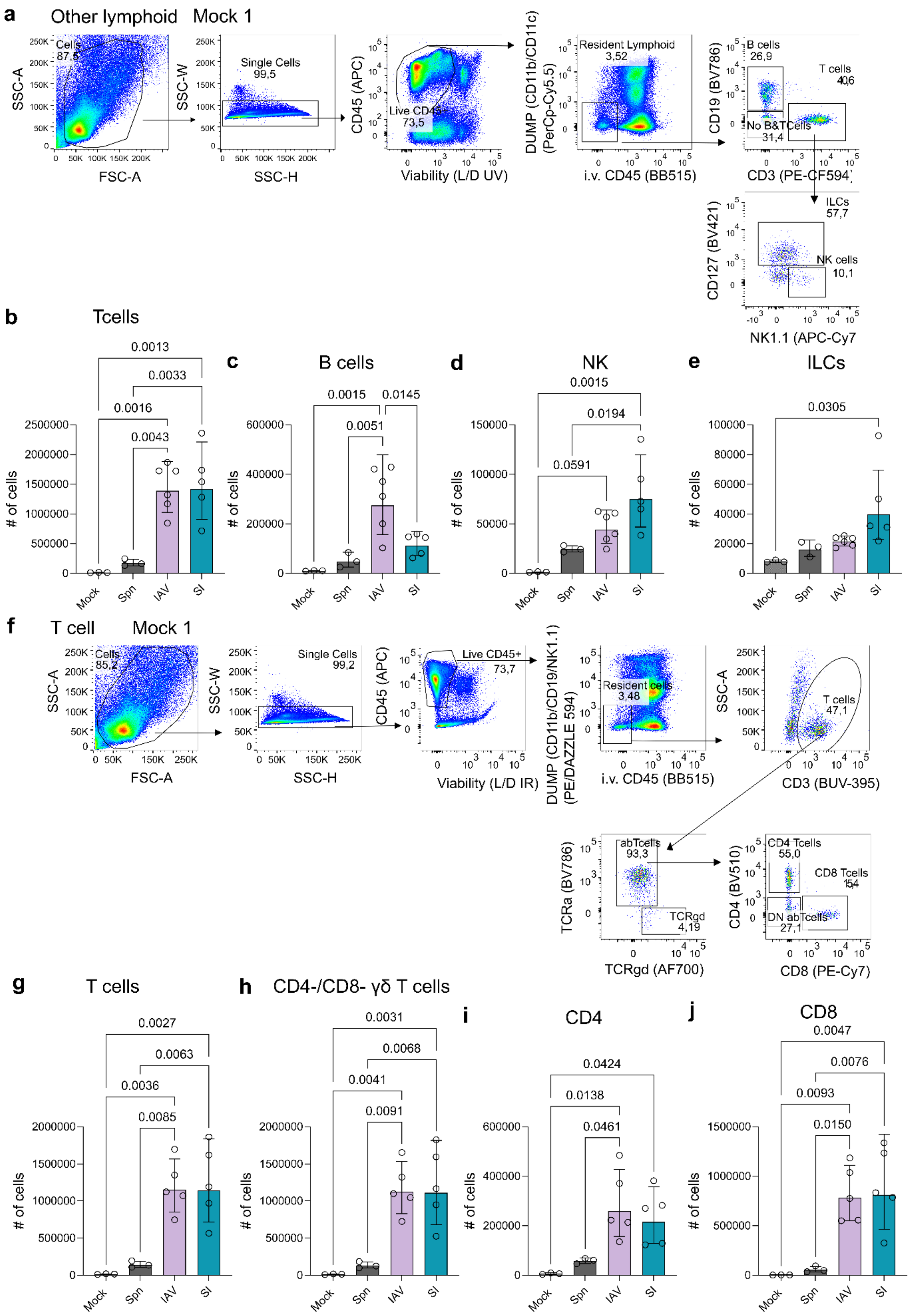

a
Other lymphoid Mock 1
Cells 87,5
SSC-A
FSC-A
Single Cells 99,5
SSC-W
SSC-H
Live CD45+ 73,5
CD45 (APC)
Viability (L/D UV)
Resident Lymphoid 3,52
DUMP (CD11b/CD11c) (PerCp-Cy5.5)
i.v. CD45 (BB515)
B cells 26,9
T cells 40,6
No B&TCells 31,4
CD19 (BV786)
CD3 (PE-CF594)
ILCs 57,7
NK cells 10,1
CD127 (BV421)
NK1.1 (APC-Cy7
b
Tcells
0.0013
0.0033
0.0016
0.0043
### of cells
Mock
Spn
IAV
SI
c
B cells
0.0015
0.0145
0.0051
d
NK
0.0015
0.0194
0.0591
e
ILCs
0.0305
f
T cell Mock 1
Cells 85,2
SSC-A
FSC-A
Single Cells 99,2
SSC-W
SSC-H
Live CD45+ 73,7
CD45 (APC)
Viability (L/D IR)
Resident cells 3,48
DUMP (CD11b/CD19/NK1.1) (PE/DAZZLE 594)
i.v. CD45 (BB515)
T cells 47,1
SSC-A
CD3 (BUV-395)
abTcells 93,3
TCRgd 4,19
TCRa (BV786)
TCRgd (AF700)
CD4 Tcells 55,0
CD8 Tcells 15,4
DN abTcells 27,1
CD4 (BV510)
CD8 (PE-Cy7)
g
T cells
0.0027
0.0063
0.0036
0.0085
h
CD4-/CD8- γδ T cells
0.0031
0.0068
0.0041
0.0091
i
CD4
0.0424
0.0138
0.0461
j
CD8
0.0047
0.0076
0.0093
0.0150

**Extended data figure 6: Influenza infection induces lymphoid cell infiltration in lung at day 8 post-infection**

**a**, Gating strategy of other lymphoid cells. **b-e**, Number of cells in lungs of Mock, IAV, Spn and SI infected mice. **(b)** T cells, **(c)** B cells, **(d)** Natural Killer, **(e)** Innate lymphoid cells, **f,** Gating strategy of T cells. **g-j,** Number of cells in lungs of Mock, IAV, Spn and SI infected mice. **(g)** T cells, **(h)** γδ T cells, **(i)** T cells CD4+ and **(j)** T cell CD8+ cells (n = 3-6 per group). Data are shown as geometric means ± geometric s.d. Statistical significance was determined by ordinary one-way ANOVA.

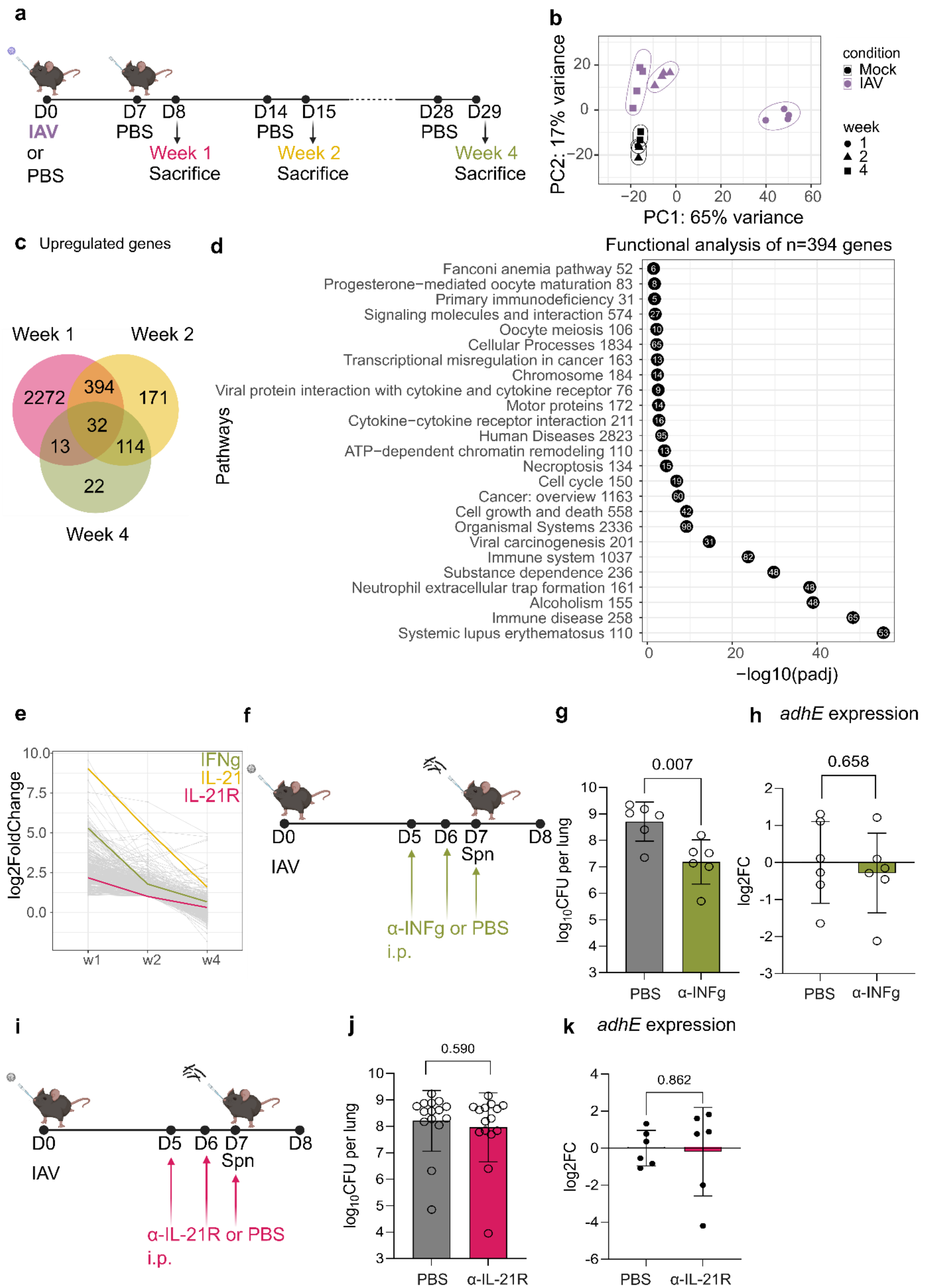
a
D0
IAV
or
PBS
D7
PBS
D8
Week 1
Sacrifice
D14
PBS
D15
Week 2
Sacrifice
D28
PBS
D29
Week 4
Sacrifice
b
PC2: 17% variance
PC1: 65% variance
condition
Mock
IAV
week
1
2
4
c
Upregulated genes
Week 1
Week 2
Week 4
2272
394
171
32
13
114
22
d
Functional analysis of n=394 genes
Pathways
Fanconi anemia pathway 52
Progesterone−mediated oocyte maturation 83
Primary immunodeficiency 31
Signaling molecules and interaction 574
Oocyte meiosis 106
Cellular Processes 1834
Transcriptional misregulation in cancer 163
Chromosome 184
Viral protein interaction with cytokine and cytokine receptor 76
Motor proteins 172
Cytokine−cytokine receptor interaction 211
Human Diseases 2823
ATP−dependent chromatin remodeling 110
Necroptosis 134
Cell cycle 150
Cancer: overview 1163
Cell growth and death 558
Organismal Systems 2336
Viral carcinogenesis 201
Immune system 1037
Substance dependence 236
Neutrophil extracellular trap formation 161
Alcoholism 155
Immune disease 258
Systemic lupus erythematosus 110
−log10(padj)
e
log2FoldChange
IFNg
IL-21
IL-21R
w1
w2
w4
f
D0
IAV
D5
D6
D7
Spn
D8
α-INFg or PBS
i.p.
g
0.007
log10CFU per lung
PBS
α-INFg
h
adhE expression
0.658
log2FC
PBS
α-INFg
i
D0
IAV
D5
D6
D7
Spn
D8
α-IL-21R or PBS
i.p.
j
0.590
log10CFU per lung
PBS
α-IL-21R
k
adhE expression
0.862
log2FC
PBS
α-IL-21R

**Extended data figure 7: Host transcriptional adaptations to IAV infection**

**a**, Schematic of the mouse infection model. Mice were intranasally (i.n.) primed with PBS or influenza A virus (IAV; 10 PFU) and subsequently administered PBS at 7, 14 or 28 days post IAV infection. Mice were euthanized 24 h after PBS intranasal administration. **b**, Principal component analysis (PCA) of host transcriptomes recovered from Mock and IAV infected mice at one week (circles), two weeks (triangles) and four weeks (squares) following IAV infection (n = 2-5 per group). **c**, Venn diagram showing host genes upregulated at one week, two weeks and four weeks following IAV infection. **d**, Pathway enrichment analysis of 394 genes commonly upregulated at weeks 1 and 2, but not at week 4, following IAV infection. Enrichment was assessed using a hypergeometric test. Numbers at the end of each pathway indicate the total number of genes annotated to that pathway, whereas numbers within circles denote the number of genes from the dataset. **e**, $Log_2$ fold change of the 394 genes upregulated at weeks 1 and 2, but no longer upregulated at week 4. IFNG (green), IL21 (yellow) and IL21R (pink) are highlighted. **f**, Schematic of the mouse infection model. Mice were primed with IAV at day 0, followed by intraperitoneal administration of anti-IFN-g antibody (α-IFN-g) or PBS at days 5, 6 and 7 post IAV infection. Mice were subsequently infected with *S. pneumoniae* at day 7. **g**, Lung bacterial burdens in superinfected mice and treated with PBS or anti-IFN-g (n = 6 per group; one experiment; geometric means ± geometric s.d. are shown). **h**, *S. pneumoniae adhE* expression in superinfected mice treated with PBS or anti-IFN-g (n = 6; one experiment; means ± s.d. are shown). **i**, Schematic of the mouse infection model. Mice were primed with IAV at day 0, followed by intraperitoneal administration of anti-IL-21R antibody (α-IL-21R) or PBS at days 5, 6 and 7 post IAV infection. Mice were subsequently infected with *S. pneumoniae* at day 7. **j**, Lung bacterial burdens in superinfected mice treated with PBS or anti-IL-21R (n = 15 per group; data pooled from two independent experiments; geometric means ± geometric s.d. are shown). **h**, *S. pneumoniae adhE* expression in superinfected mice treated with PBS or anti-IL-21R (n = 5-6; one experiment; means ± s.d. are shown). Statistical significance was determined by two-tailed unpaired *t*-tests **(g,h,j,k)**.

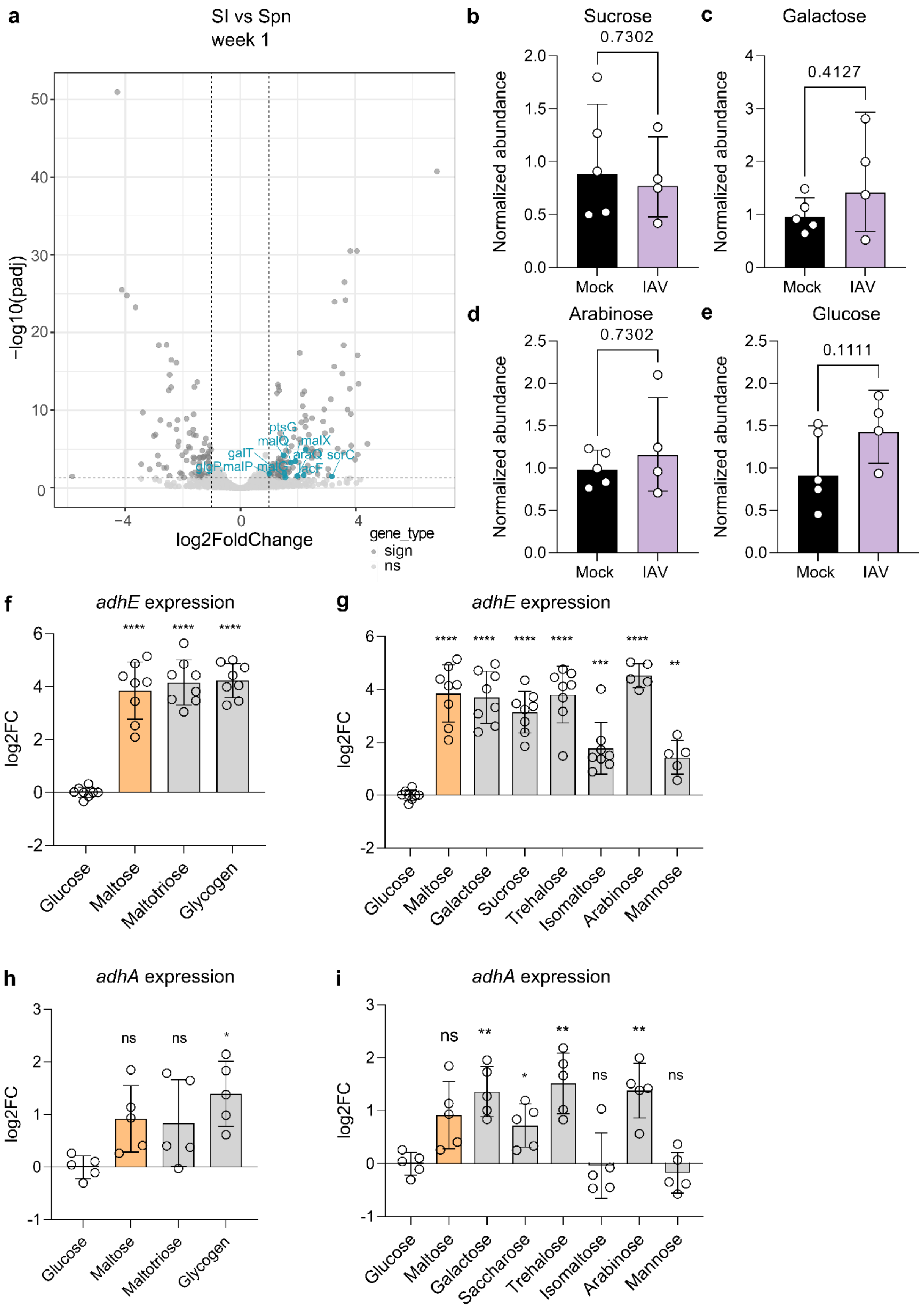

a
SI vs Spn
week 1
−log10(padj)
log2FoldChange
gene_type
sign
ns
ptsG
malQ
malX
galT
araQ
sorC
glgP
malP
lacF
b
Sucrose
0.7302
c
Galactose
0.4127
d
Arabinose
0.7302
e
Glucose
0.1111
Normalized abundance
Mock
IAV
f
adhE expression
g
adhE expression
log2FC
Glucose
Maltose
Maltotriose
Glycogen
Galactose
Sucrose
Trehalose
Isomaltose
Arabinose
Mannose
h
adhA expression
i
adhA expression
Saccharose
ns

**Extended data figure 8: Alternative carbohydrates induced by IAV infection trigger adh upregulation**

**a**, Volcano plots showing differentially expressed *S. pneumoniae* genes during SI compared with Spn infection at week 1. Genes upregulated that are related to carbohydrate uptake and metabolism are indicated by blue dots. **b-e**, **(b)** Sucrose, galactose **(c)**, arabinose **(d)** and glucose **(e)** levels in cells from bronchoalveolar lavage fluid from mock- or IAV-infected mice, measured by targeted metabolomic analysis using GC–MS. Metabolite levels are shown as relative abundance normalized to mock-infected samples and to total sample signal. Dots indicate individual mice, bars depict geometric means ± geometric s.d. (n = 4-5; one representative of at least two experiments shown). **f**, *adhE* expression in *S. pneumoniae* cultured in chemically defined medium supplemented with 1% of glucose, maltose, maltotriose and glycogen for 30 min. Dots represent individual bacterial cultures (n = 8, data pooled from at least two independent experiments; means ± s.d. are shown). **g**, *adhE* expression in *S. pneumoniae* cultured in chemically defined medium supplemented with 1% of glucose, maltose, galactose, sucrose, trehalose, isomaltose, arabinose and mannose for 30 min. Dots represent individual bacterial cultures (n = 5-8, data pooled from at least two independent experiments; means ± s.d. are shown). **h**, *adhA* expression in *S. pneumoniae* cultured in chemically defined medium supplemented with 1% of glucose, maltose, maltotriose and glycogen for 30 min. Dots represent individual bacterial cultures (n = 5-8, data pooled from at least two independent experiments; means ± s.d. are shown). **i,** *adhA* expression in *S. pneumoniae* cultured in chemically defined medium supplemented with 1% of glucose, maltose, galactose, sucrose, trehalose, isomaltose, arabinose and mannose for 30 min. Dots represent individual bacterial cultures (n = 5, data pooled from at least two independent experiments; means ± s.d. are shown). Statistical significance was determined by unpaired two-tailed Mann–Whitney tests **(b-e)** and two-tailed unpaired *t*-tests **(f- i).**

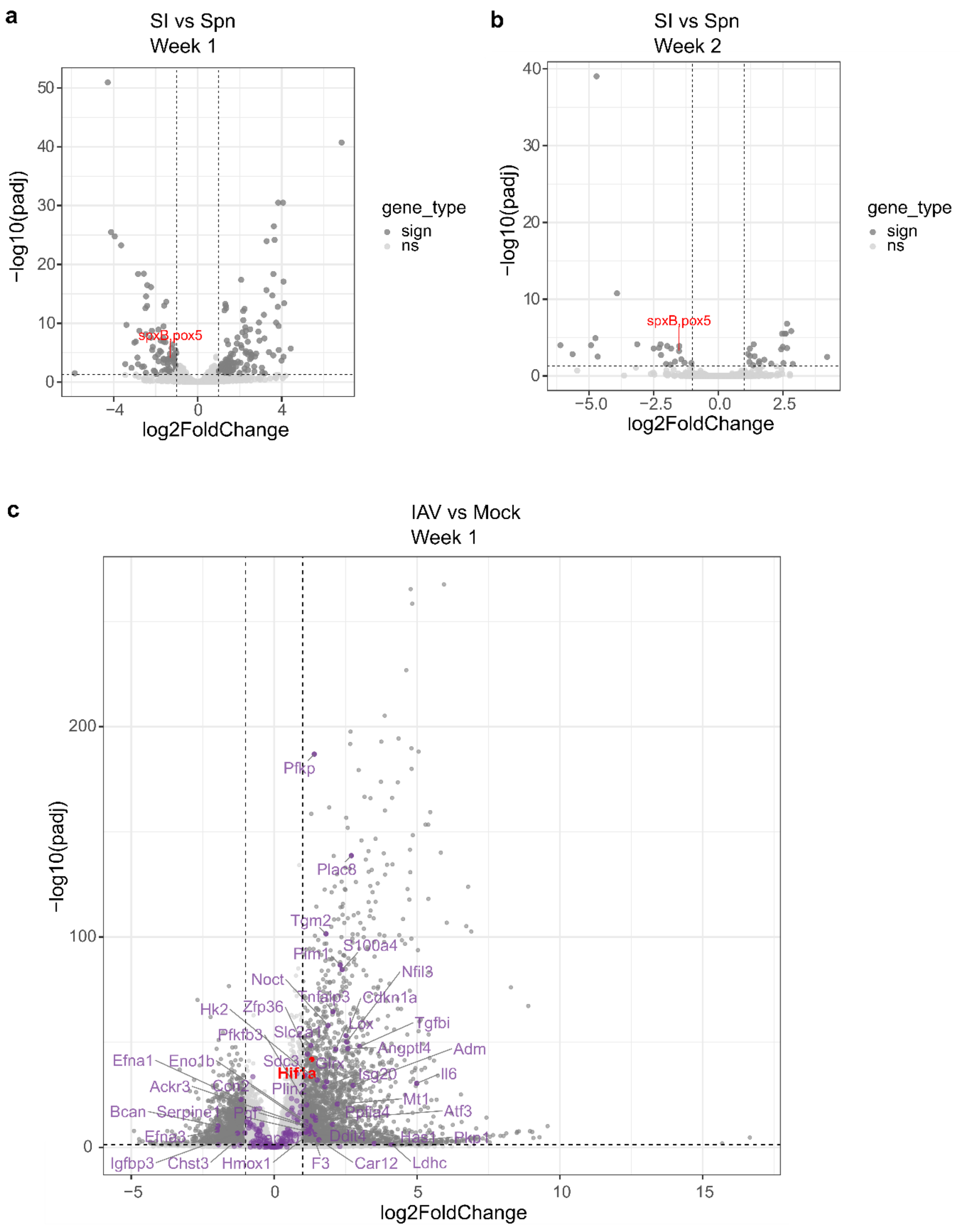

a
SI vs Spn
Week 1
-log10(padj)
log2FoldChange
spxB,pox5
gene_type
sign
ns
b
SI vs Spn
Week 2
-log10(padj)
log2FoldChange
spxB,pox5
gene_type
sign
ns
c
IAV vs Mock
Week 1
-log10(padj)
log2FoldChange
Pfkp
Plac8
Tgm2
Plin1
S100a4
Noct
Nfil3
Tnfaip3
Cdkn1a
Hk2
Zfp36
Lox
Tgfbi
Pfkfb3
Slc2a1
Angptl4
Adm
Efna1
Eno1b
Sdc3
Hif1a
Isg20
Il6
Ackr3
Ccn2
Plin2
Mt1
Bcan
Serpine1
Pgf
Atf3
Efna3
Ddit4
Has1
Igfbp3
Chst3
Hmox1
F3
Car12
Ldhc

**Extended data figure 9: Hypoxia-related genes are differentially expressed in host and bacterial transcriptomes**

**a,b**, Volcano plots showing differentially expressed *S. pneumoniae* genes during SI compared with Spn infection at (a) one week and (b) two weeks following IAV infection. *spxB* is indicated in red. **c,** Volcano plot of host transcriptional response at one week following IAV infection. Hypoxia hallmark from MSigDB are indicated in purple and *HiF1a* in red. Data are representative of at least two experiments.

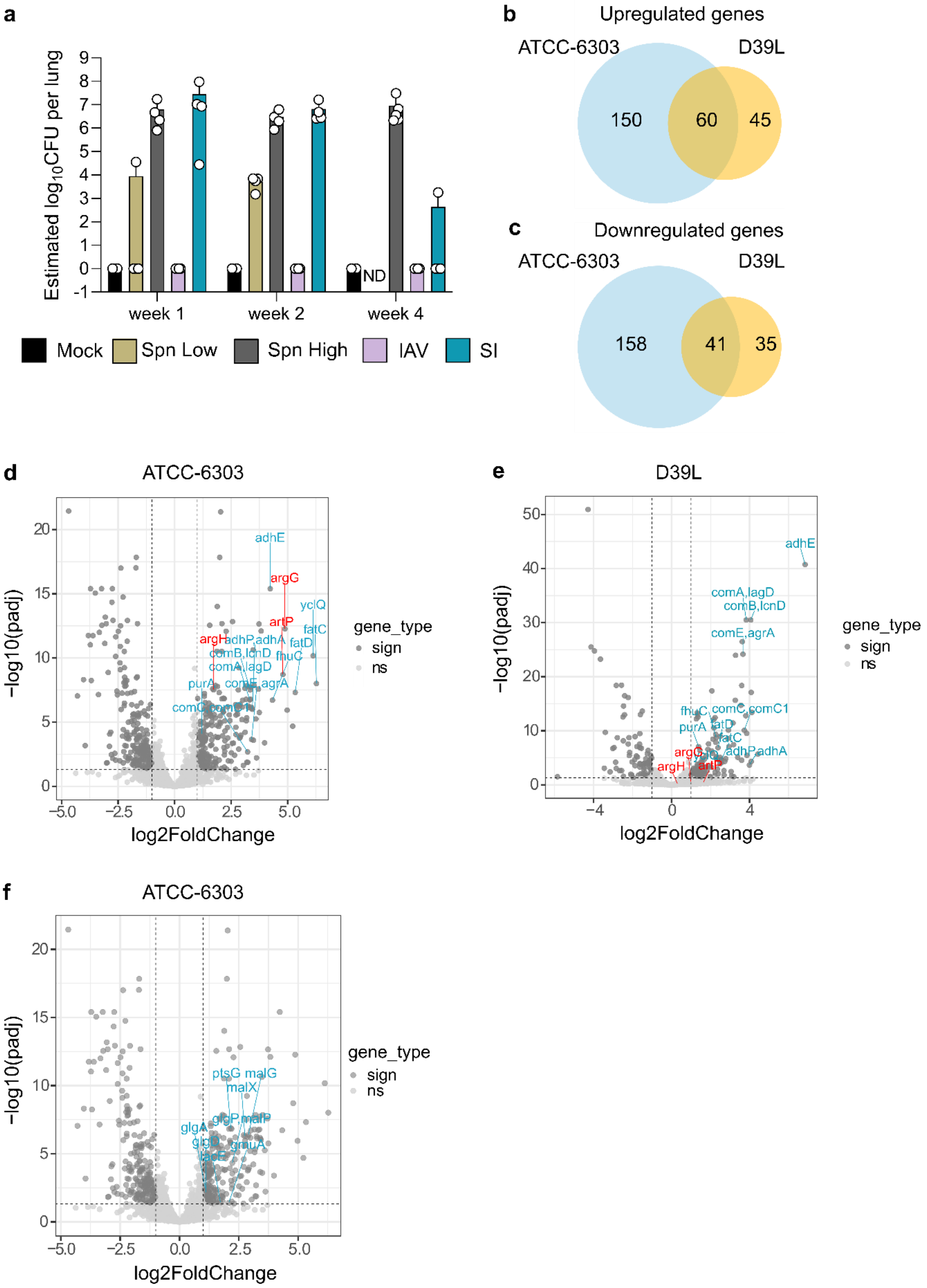
a
Estimated log10CFU per lung
9
8
7
6
5
4
3
2
1
0
-1
ND
week 1
week 2
week 4
Mock
Spn Low
Spn High
IAV
SI
b
Upregulated genes
ATCC-6303
D39L
150
60
45
c
Downregulated genes
ATCC-6303
D39L
158
41
35
d
ATCC-6303
−log10(padj)
log2FoldChange
adhE
argG
yclQ
artP
fatC
argH
adhP,adhA
fatD
comB,lcnD
fhuC
comA,lagD
purA
comE,agrA
comC,comC1
gene_type
sign
ns
e
D39L
−log10(padj)
log2FoldChange
adhE
comA,lagD
comB,lcnD
comE,agrA
fhuC
comC,comC1
purA
fatD
fatC
argG
adhP,adhA
argH
artP
gene_type
sign
ns
f
ATCC-6303
−log10(padj)
log2FoldChange
ptsG
malG
malX
glgA
glgP
malP
glgD
gmuA
lacE
gene_type
sign
ns

**Extended data figure 10: Comparison of *S. pneumoniae* serotype 2 and serotype 3 adaptation to superinfection**

**a**, Estimation of *S. pneumoniae* ATCC-6303 titers at one, two and four weeks following IAV infection, based on expression of the housekeeping gene *gyrB,* as determined by RT-qPCR. Dots indicate individual mice (n = 2-5 per group; geometric means ± geometric s.d. are shown).

**b,c**, Venn diagram showing *S. pneumoniae* genes **(b)** upregulated and **(c)** downregulated that are shared by D39L and ATCC-6303 during superinfection at week 1.

**d,e**, Volcano plots showing differentially expressed genes during SI compared with Spn infection at week 1 for *S. pneumoniae* **(d)** ATCC-6303 and **(e)** D39L. Common upregulated genes are shown in blue and strain-specific upregulated genes in red.

**f**, Volcano plot showing differentially expressed *S. pneumoniae* ATCC-6303 genes during SI compared with Spn infection at week 1. Genes upregulated that are related to carbohydrate uptake and metabolism are indicated by blue dots.

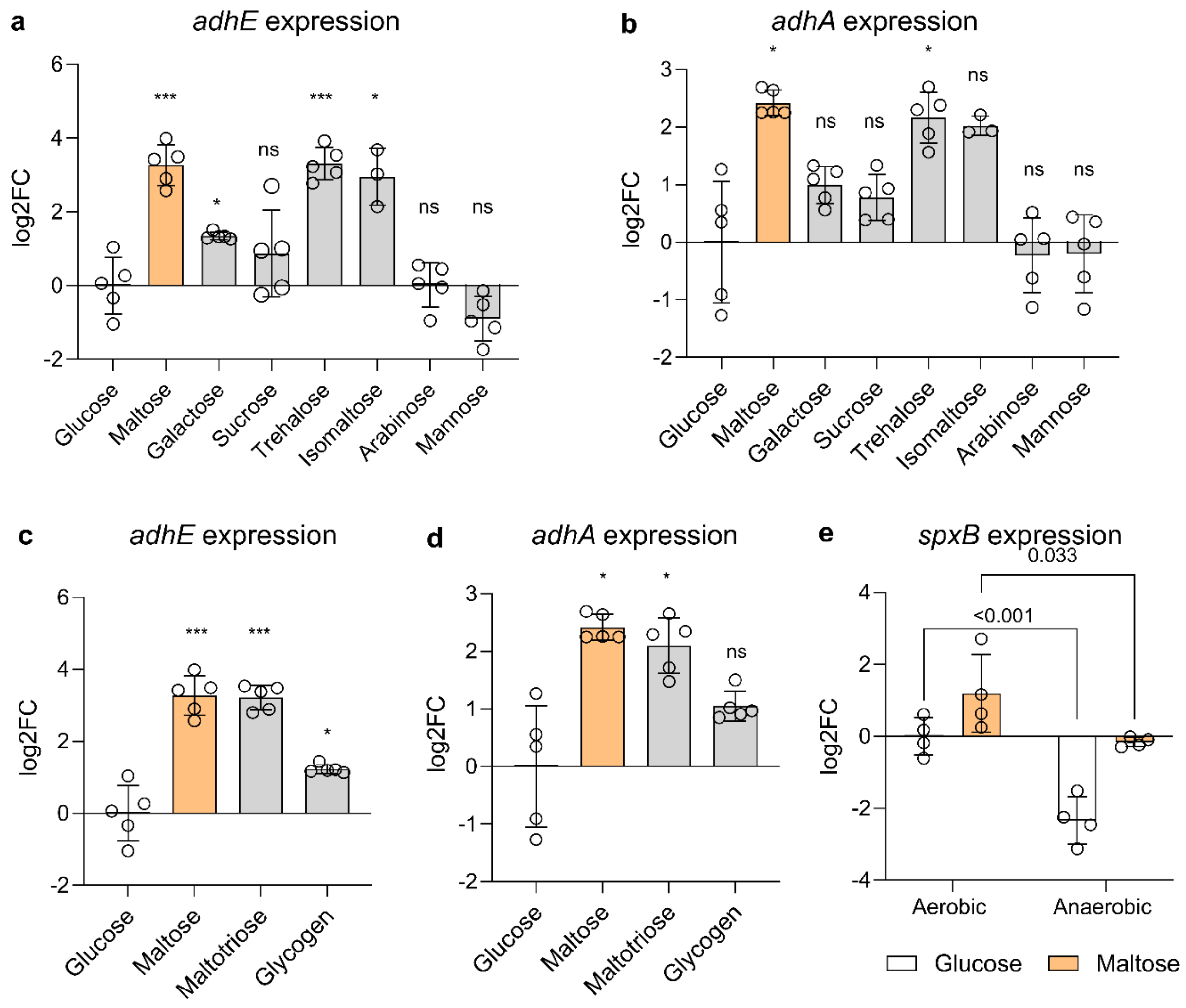

a
adhE expression
log2FC
6
4
2
0
-2
***
*
ns
***
*
ns
ns
Glucose
Maltose
Galactose
Sucrose
Trehalose
Isomaltose
Arabinose
Mannose
b
adhA expression
log2FC
3
2
1
0
-1
-2
*
ns
ns
*
ns
ns
ns
Glucose
Maltose
Galactose
Sucrose
Trehalose
Isomaltose
Arabinose
Mannose
c
adhE expression
log2FC
6
4
2
0
-2
***
***
*
Glucose
Maltose
Maltotriose
Glycogen
d
adhA expression
log2FC
3
2
1
0
-1
-2
*
*
ns
Glucose
Maltose
Maltotriose
Glycogen
e
spxB expression
0.033
<0.001
log2FC
4
2
0
-2
-4
Aerobic
Anaerobic
Glucose
Maltose

**Extended data figure 11: *S. pneumoniae* serotype 3 (ATCC-6303) adaptation to alternative carbon sources and oxygen availability**

**a,b**, *adhE* **(a)** and *adhA* **(b)** expression in *S. pneumoniae* cultured in chemically defined medium supplemented with 1% of glucose, maltose, galactose, sucrose, trehalose, isomaltose, arabinose or mannose. Dots represent individual bacterial cultures (n = 3-5, data pooled from at least two independent experiments; means ± s.d. are shown). **c,d**, *adhE* **(c)** and *adhA* **(d)** expression in *S. pneumoniae* cultured in chemically defined medium supplemented with 1% of glucose, maltose, maltotriose or glycogen for 30 min. Dots represent individual bacterial cultures (n = 5, data pooled from at least two independent experiments; means ± s.d. are shown). **e**, *spxB* expression in *S. pneumoniae* cultured in chemically defined medium supplemented with glucose or maltose under aerobic or anaerobic conditions for 30 min. Dots represent individual bacterial cultures (n = 4, data pooled from at least two independent experiments; means ± s.d. are shown). Statistical significance was determined by two-tailed unpaired *t*-tests **(a-e)**.
